# Electrophysiological measures from human iPSC-derived neurons are associated with schizophrenia clinical status and predict individual cognitive performance

**DOI:** 10.1101/2021.04.08.437289

**Authors:** Stephanie Cerceo Page, Srinidhi Rao Sripathy, Federica Farinelli, Zengyou Ye, Yanhong Wang, Daniel J Hiler, Elizabeth A Pattie, Claudia V Nguyen, Madhavi Tippani, Rebecca L. Moses, Huei-Ying Chen, Matthew Nguyen Tran, Nicholas J Eagles, Joshua M Stolz, Joseph L Catallini, Olivia R Soudry, Dwight Dickinson, Karen F Berman, Jose A Apud, Daniel R Weinberger, Keri Martinowich, Andrew E Jaffe, Richard E Straub, Brady J Maher

## Abstract

Neurons derived from human induced pluripotent stem cells (hiPSCs) have been used to model basic cellular aspects of neuropsychiatric disorders, but the relationship between the emergent phenotypes and the clinical characteristics of donor individuals has been unclear. We analyzed RNA expression and indices of cellular function in hiPSC-derived neural progenitors and cortical neurons generated from 13 individuals with high polygenic risk scores (PRS) for schizophrenia and a clinical diagnosis of schizophrenia, along with 15 neurotypical individuals with low PRS. We identified electrophysiological measures associated with diagnosis that implicated altered Na^+^ channel function and GABA-ergic neurotransmission. Importantly, electrophysiological measures predicted cardinal clinical and cognitive features found in these schizophrenia patients. The identification of basic neuronal physiological properties related to core clinical characteristics of illness is a potentially critical step in generating leads for novel therapeutics.

## Introduction

Schizophrenia (SCZ) is a complex, highly heritable, “polygenic” disorder with considerable cost to affected individuals, their families and society^1^. Marked clinical heterogeneity supports the idea that SCZ is a spectrum of disorders with overlapping symptomatology, rather than a single disease. SCZ is characterized by positive symptoms including hallucinations and delusions, negative symptoms such as social and emotional withdrawal, and cognitive symptoms including deficits in executive function and decision making. Negative symptoms^2^ and cognitive deficits^3^ in SCZ are still only poorly addressed by current treatments, all of which have side effects that often result in poor medication adherence. Although SCZ is not typically diagnosed until early adulthood, versions of a “neurodevelopmental hypothesis”^4^ propose that altered signaling during early brain development leads to altered connectivity/function of neural circuits in later critical maturational periods, and eventually leads to onset of clinical symptoms and cognitive deficits.

Substantial genetic and environmental complexity and heterogeneity are hallmarks of common, polygenic disorders, posing serious challenges to the mechanistic understanding necessary for advances in drug design. Genome-wide association studies (GWAS) have implicated at least 270 risk loci for SCZ, with each risk-associated variant tagging multiple haplotypes containing functional variants that confer only a small degree of risk^5^, and it is predicted, at least at the population level, that there exists thousands of such risk-related genes. Derived from GWAS, the polygenic risk score (PRS)^6, 7^ is a relative, cumulative index of genomic risk for disease, calculated for each individual based on independent genetic risk markers. PRS is based on common genetic variation, and has become an important tool in SCZ research that has the potential to identify undiagnosed individuals for whom the risk of developing SCZ is relatively high, as well as in choosing subjects for molecular and clinical studies. However, very little is known about how these common variants and risk genes interact with each other and with the environment during neurodevelopment to initiate pathophysiology. Identification of simpler and more tractable, genome-type/cellular phenotype relationships may facilitate progress in discovering biological mechanisms by which genomic risk for SCZ translates into more complex phenotypic states, potentially including diagnostic symptoms and cognitive deficits.

The advent of human induced pluripotent stem cell (hiPSC) technology to model biological aspects of neuropsychiatric disease^8^ has enabled specific and detailed characterization and manipulation of human neural cells *in vitro*, without the limitations inherent in animal models and postmortem human brain studies. Several previous studies have used hiPSC technologies to model biological aspects of SCZ in a variety of downstream cell types including neural progenitors, glial cells, and both excitatory and inhibitory neurons^9–18^. Although PRS was not used as a criteria for sample selection in those studies, associations between cellular phenotypes and SCZ diagnosis were observed, suggesting that the hiPSC model has considerable potential to identify biological dysfunction that might contribute to disease risk. However, a major uncertainty that remains in all published studies is whether any of the observed cellular phenotypes have any meaningful relationship to specific clinical or cognitive characteristics of the individual donors. This is an especially critical consideration in efforts to validate potential cell models of neuropsychiatric disorders because such illnesses do not have pathognomonic cellular pathology.

In this study we investigated the cellular impact of common genetic risk for SCZ to better understand its role in early neural development, and to possibly identify cellular phenotypes that might relate to cardinal clinical and cognitive features of the adult donors. Because the vast majority of cases of SCZ are caused by common variant risk, we focused on PRS to contrast high genetic risk patients compared with neurotypicals of low genetic risk. Based on PRS, we selected patient fibroblast lines from the participants in the NIMH Clinical Brain Disorders Branch Sibling Study of Schizophrenia (D.R. Weinberger, PI), which captured extensive clinical and cognitive data on each subject. Here, we describe the design, execution, and results of our investigation into the relationships between the molecular, cellular, and physiological properties of hiPSC-derived neural progenitor cells (NPCs) and cortical neurons with clinical status, clinical symptoms, and cognitive performance. Importantly, we look beyond case-control differences, and identify within-case patterns of association that could serve to illuminate dimensions of illness and subgroups of patients that may be suitable for targeted treatments. The results demonstrate for the first time that common genetic variants associated with SCZ converge on select electrophysiological mechanisms that relate to clinically relevant features of SCZ, underscoring the potential of the findings for biomarker identification and perhaps downstream drug development.

## Results

### Reprogramming and characterization of hiPSC lines

The study included 13 high polygenic risk score (PRS) individuals with SCZ (16 total cell lines: 2 lines each for 3 donors and 1 line each for 10 donors) and 15 low PRS neurotypical (CON) (20 total cell lines: 2 lines each for 5 donors and 1 each for 10 donors) (Fig. 1a **and Supplementary Table 1**). These hiPSC lines were generated as previously described^19^, and no differences in pluripotency were observed between SCZ and CON lines (Fig. 1b,c, Extended Data Fig. 1). The hiPSCs were first differentiated into SOX2-and PAX6-expressing NPCs, and then further differentiated into forebrain-specific cortical neurons, which were co-cultured on rat astrocytes as previously described (Fig. 1d,e)^9, 12^.

**Figure 1.**
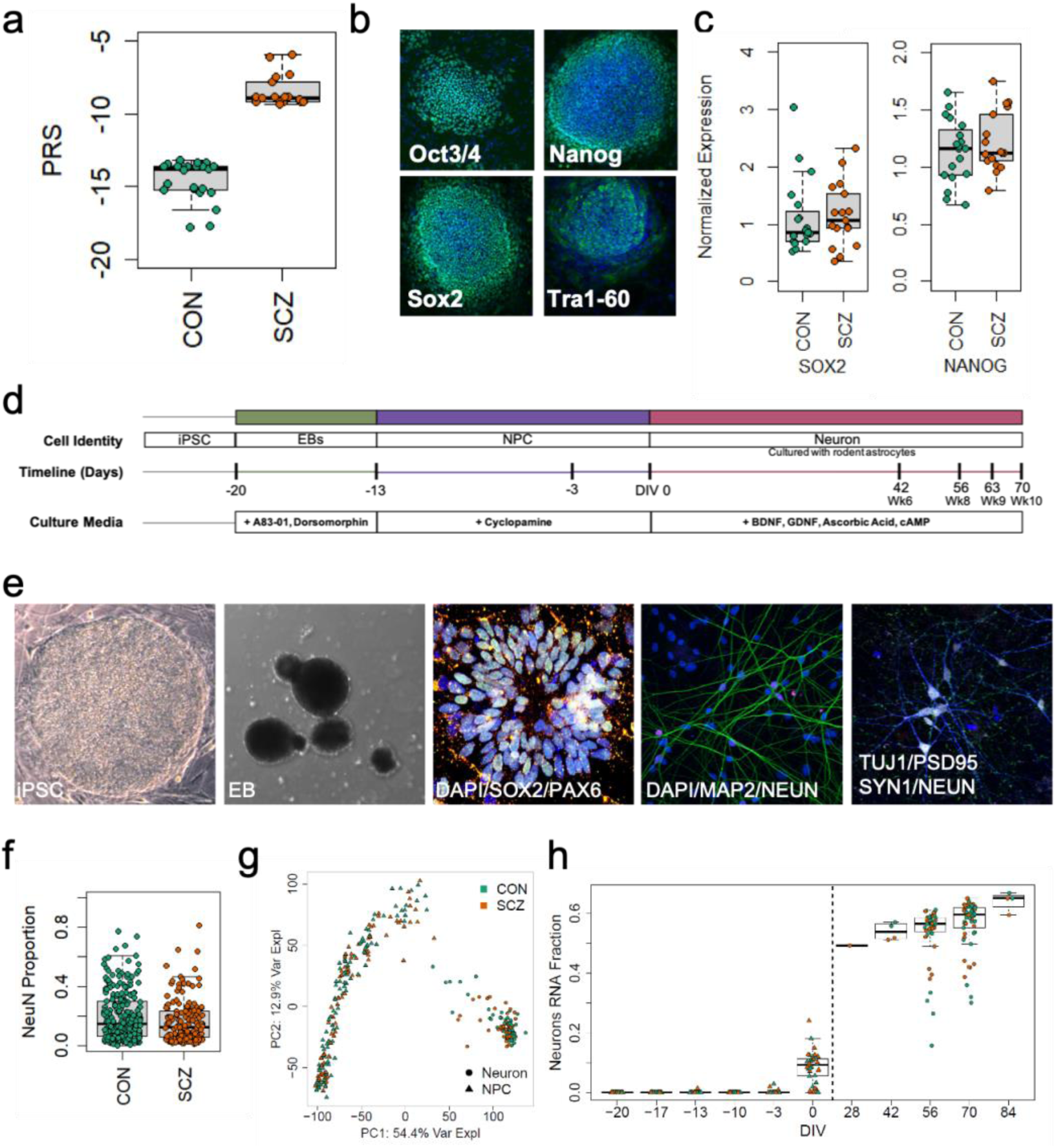
Description of experimental lines and experiment overview. **a,** Polygenic risk scores (PRS) of neurotypical controls (CON) and individuals with schizophrenia (SCZ) included in this study (CON=-14.829 ± 1.535, N=15; SCZ=-8.375 ± 1.1736, N=13; p= 1.78E-12). **b,** Pluripotency of reprogrammed lines was verified by identification of NANOG+, SOX2+, OCT3/4+, and TRA-1-60+ colonies colocalized with DAPI using immunocytochemistry. **c,** qPCR quantification of stem cell markers *SOX2* (CON=1.152 ± 0.151, SCZ=1.162 ± 0.133, p=0.96, N=35 from 28 genomes) and *NANOG* (CON=1.133 ± 0.062, SCZ=1.222 ± 0.063, p=0.32, N=35 from 28 genomes), normalized to embryonic stem cell line H1. **d,** hiPSCs were differentiated into neurons in multiple batches. **e,** Representative images of induced pluripotent stem cell colony (hiPSC, brightfield), embryoid body (EB, brightfield), immunofluorescence of neural rosette at two days, DAPI (blue), SOX2 (green), PAX6 (red), immunofluorescence of neurons at DIV56, DAPI (blue), MAP2 (green), NeuN (red), immunofluorescence of neurons showing synaptic puncta at DIV70, Tuj1 (blue), PSD95 (green), SYNAPSIN1 (red), NeuN (white). **f,** Quantification of the percentage of neurons by diagnosis (CON=0.186 ± 0.01, N=229 cultures from 18 lines; SCZ=0.167 ± 0.012, p=0.43, N=156 cultures from 14 lines). **g,** PCA of RNA-seq at multiple time points during culture differentiation (hiPSC, NPC, DIV56 and DIV70 neurons) demonstrates that the largest source of variation (top principal component, PC1, representing 54.4% of the variance explained) corresponds to cell state. **h,** Fraction of RNA attributed to neurons increases across differentiation stages in CON and SCZ lines.

### Generation of cortical neurons and molecular profiling

For all experiments, differentiation and data collection were performed blinded to diagnosis. We quantified the cellular composition of cultures across 4 rounds of differentiation and observed no differences in number of neurons (Fig. 1f) or neuronal types by SCZ (Extended Data Fig. 2a-c) or by line (Extended Data Fig. 3). Additionally, gene expression corresponding to neuron identity and maturity was assessed by RNA sequencing^9^ where the largest source of variation (54.4%) corresponded to cell state (Fig. 1g, Extended Data Fig. 4a). The fraction of RNA from neurons increased over time (Fig. 1h), while the hiPSC fraction declined over time (Extended Data Fig. 4c) and NPC fraction peaked at “days *in vitro*” 0 (DIV0; Extended Data Fig. 4d).

We performed linear mixed effects modeling on 23,119 expressed genes assessing for a main effect of SCZ^20^, adjusting for fixed effects corresponding to cellular maturity, human alignment fraction, human gene assignment rate, and DIV of the neurons (binary, corresponding to 8 or 10 weeks), and treating each donor as a random intercept to account for repeated measures. We identified 68 differentially expressed genes (DEGs) at a false discovery rate (FDR)<0.1, with median absolute fold changes of 1.8 (interquartile range: 1.34-2.45, Extended Data Fig. 5a). We verified that no differentially expressed genes for SCZ were found in the rat genome - corresponding to effects in astrocytes - at an FDR<0.99 using an analogous statistical model. We then performed gene set enrichment.

Using a larger set of DEGs (at p<0.005) and predefined Gene Ontology sets, we identified pathways related to WNT signaling and forebrain development (Extended Data Fig. 5b). We further found that SCZ DEGs correlated across cell states, such that DEGs in NPCs were preserved as DEGs in neurons. This developmental overlap in SCZ DEGs suggests that expression differences found in our neurons started earlier in development/differentiation, and that for some genes, differences in NPCs may predict differences in neurons (Extended Data Fig. 5c-e)^10^. Finally, we compared DEGs in our hiPSC-derived neurons to SCZ state-related DEG and risk-related TWAS in the post-mortem dorsolateral prefrontal cortex (DLPFC; Extended Data Fig. 5f,g), and similar to a previous study^21^, we found no association between SCZ illness state (DEGs) and SCZ genetic risk (TWAS). Together, these results indicate that diagnosis or DEG associated with diagnosis do not significantly impact the generation of subtypes of cortical forebrain neurons thus enabling downstream cellular and physiological comparisons between genomes and diagnosis.

### Initial physiological characterization

We assessed cortical neuron function using Ca^2+^ imaging and whole-cell electrophysiology. Our differentiation protocol consistently generated electrically active neurons that displayed spontaneous and synchronous network activity (Extended Data Fig. 6a-g). We quantified Ca^2+^ transients at DIV42 and DIV63 from 13,602 individual neurons across 871 videos, and observed no effect of diagnosis on the proportion of active cells (Extended Data Fig. 6a-c, Extended Data Fig. 7a), the frequency of events (Extended Data Fig. 7b,c), or the synchronicity of events (Extended Data Fig. 7d-f). Whole-cell recordings from over 1,000 neurons at DIV56 and DIV70 from multiple differentiation runs confirmed appropriate developmental maturation of intrinsic membrane properties (Extended Data Fig. 8), the ability to fire trains of action potentials (AP; **Supplementary Table 2**), and spontaneous synaptic transmission (Extended Data Fig. 5e-g), collectively indicating that our directed differentiation protocol generates cortical neurons that display intrinsic electrical activity and synaptic connectivity that results in functional neuronal networks.

### Analysis of association between electrophysiological measures and diagnosis

Three electrophysiological measures related to Na^+^ channel function were associated (p<0.05) with diagnosis (Fig. 2a, **Supplementary Table 2**). Lines derived from SCZ donors (SCZ lines) showed an increase in membrane resistance (Fig. 2a, p=0.025), an increased number of Na^+^ current peaks in response to a voltage ramp protocol (p=0.038; Fig. 2a-c, Extended Data Fig. 9a,b), and a decrease in the activation threshold of the second Na^+^ peak (p=0.009; Fig. 2a). Further examination of Na^+^ currents generated during the ramp protocol showed they are TTX sensitive (Extended Data Fig. 10a,b), and that their frequency correlates with physiologically relevant indices indicative of neuronal maturation (Extended Data Fig. 10c-e). Interestingly, increased Na^+^ currents are related to more mature AP kinetics, even though SCZ lines show increased membrane resistance, which is typical of a less mature neuron that is expressing fewer ion channels on its plasma membrane. In addition, we observed that the frequency of spontaneous inhibitory synaptic currents (sIPSC) was increased in SCZ lines (p=1.12E-06, Fig. 2a,d-e, Extended Data Fig. 9e,f), while spontaneous excitatory synaptic transmission (sEPSC) was unchanged (Extended Data Fig. 6f,g, **Supplementary Table 2**), suggesting altered excitatory/inhibitory balance. The enhanced sIPSC frequency appears to be related to an increase in the number of GABA-ergic synapses and/or altered presynaptic probability of release, since the number of GABA-ergic cells (Extended Data Fig. 2), the relative expression of *SLC12A2* (protein NKCC1, p=0.66) and *SLC12A5* (protein CC2, p=0.80), and the frequency of network activity (Extended Data Fig. 7b,c) did not differ by diagnosis. Together, these results identified associations between diagnosis and specific physiological phenotypes that are related to Na^+^ channel function and GABA-ergic transmission.

**Figure 2.**
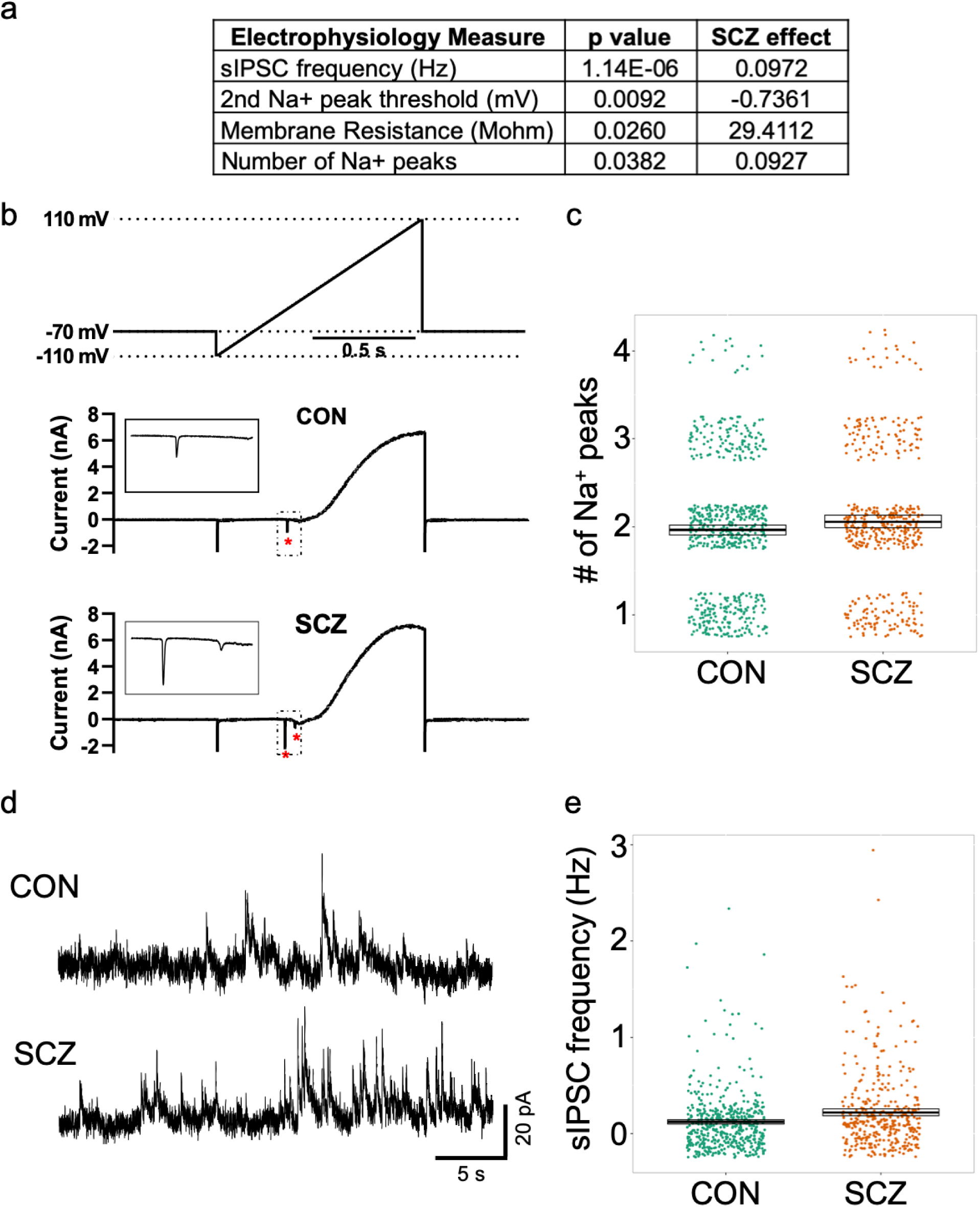
Electrophysiological differences between CON and SCZ lines. **a,** Table of electrophysiology phenotypes observed between CON and SCZ lines. **b,** Representative traces showing Na^+^ currents (* and inset) in response to a voltage ramp protocol (above). **c,** Summary data showing significantly more Na^+^ currents in SCZ neurons versus CON neurons across all genomes (SCZ effect=0.093, p=0.038, N=1074 from 28 genomes). **d,** Representative traces of spontaneous inhibitory postsynaptic currents (sIPSCs) recorded from CON and SCZ neurons at 0mV. **e,** Summary data showing significantly increased sIPSC frequency in SCZ neurons versus CON neurons across all genomes (SCZ effect=0.097, p=1.14E-6, N=903 from 28 genomes).

### Sodium channel kinetics associated with SCZ

Given that three electrophysiology measures are directly related to Na^+^ channel function (Fig 2A) and are associated with diagnosis, we performed a more detailed electrophysiological analysis of Na^+^ channel kinetics. To improve our voltage-clamp of Na^+^ currents, we used a modified internal recording solution and measured the voltage-dependence of Na^+^ channel activation and inactivation on a subset (see **Supplementary Table 1**) of SCZ (7 lines from 6 genomes) and CON (7 lines from 7 genomes) genomes (Fig. 3a,d). Using standard voltage-clamp protocols^22^, we observed that SCZ lines showed a significantly hyperpolarized shift in Na^+^ channel activation (Fig. 3b, Extended Data Fig. 11) and V1/2 activation voltage (p=0.039; Fig. 3c). In addition, SCZ lines showed a significant hyperpolarized shift in the voltage dependence of inactivation (Fig. 3e, Extended Data Fig. 11) and V1/2 inactivation voltage (p<0.0001, adjusted p<0.0035; Fig. 3f). The hyperpolarized shift in activation supports the diagnosis association with the threshold for activation of the second Na^+^ peak (Fig. 2a) and the hyperpolarized shift in inactivation supports the higher membrane resistance observed in SCZ lines (Fig. 2a), because more channels are in a nonconducting state at resting membrane potentials. Together, these results indicated that Na^+^ channel function is altered in cortical neurons derived from SCZ donors in comparison to controls. This phenotype can impact intrinsic excitability^23^ and the developmental trajectory of cortical neurons and circuits^24^, potentially representing an underlying etiological mechanism for altered connectivity implicated in schizophrenia.

**Figure 3.**
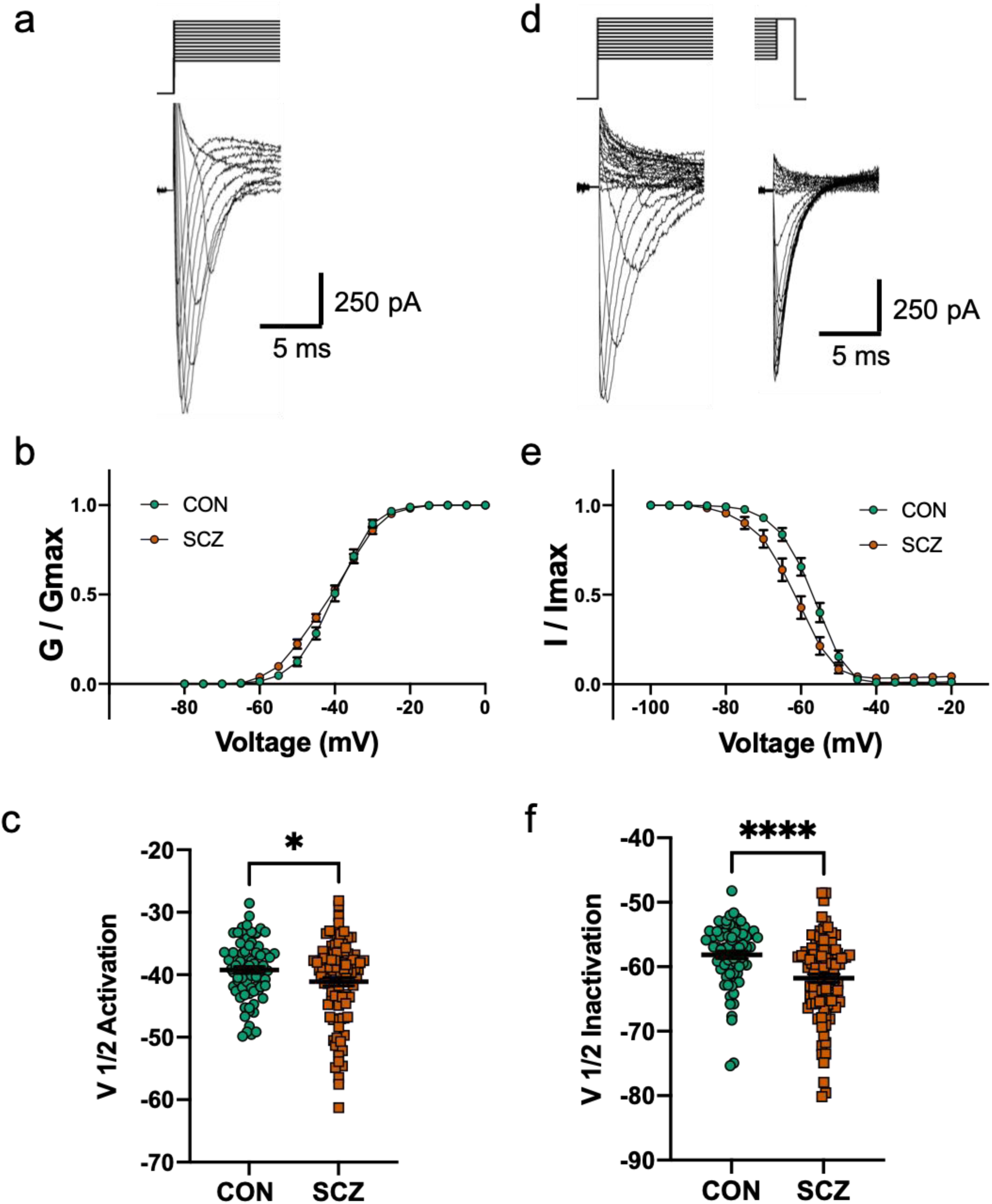
Altered Na^+^ channel activation and inactivation in 6 SCZ compared to 7 CON. **a,** Representative Na^+^ current traces in response to voltage steps (−100mV to +20mV). **b,** Voltage-dependence of steady-state activation of Na^+^ channels in SCZ and CON neurons. **c,** Summary data showing significantly hyperpolarized shift in the activation of Na^+^ channels in SCZ neurons versus CON neurons (CON, V1/2=-39.2 ± 0.6 mV; SCZ, V1/2=-41.1 ± 0.7 mV, Delta = 1.9 mV, p=0.039, N=173 from 13 genomes). **d,** Representative Na^+^ current traces in response to a steady state inactivation protocol (initial voltage steps from −130mV to 20mV; test voltage steps +20mV) to measure channel inactivation. **e,** Voltage-dependence of steady-state inactivation of Na^+^ channels in SCZ and CON neurons. **f,** Summary data showing significantly hyperpolarized shift in the inactivation of Na^+^ channels in SCZ neurons versus CON neurons (CON V1/2=-58.1 ± 0.6 mV; SCZ=-61.8 ± 0.7 mV, Delta=3.7 mV, p<0.0001, adjusted p<0.0035, N=162 from 13 genomes).

### Association of electrophysiological measures with clinical and cognitive data

Lastly, we correlated electrophysiology measures with clinical symptomatology and cognitive performance of the cell line donors. We focused on 7 electrophysiology measures that either showed association with SCZ diagnosis (Figure 2a) or were related to spontaneous synaptic transmission. Ratings for Positive and Negative Syndrome Scale items (PANSS; 7 positive, 7 negative, 16 general) captured classic diagnostic symptomatology. Regarding cognition, we tested composite factor scores for 6 cognitive domains (Verbal Memory, Nback, Visual Memory, Processing Speed, Card Sorting, and Digit Span), as well as estimated IQ and a general cognitive ability composite (“g”)^25–30^. Correlations within CON, who show limited variation on symptom ratings, but normal cognitive test score variation, were either very weak or absent (**Supplementary Table 4**).

In SCZ, however, two electrophysiological variables showed a markedly divergent pattern of associations with the clinical and cognitive variables, echoing a frequent divergence of these characteristics in patients (Figure 4a). The number of Na^+^ peaks showed directionally-consistent, positive associations with canonical positive symptom ratings (for hallucinations and delusions), whereas there were no correlations with negative symptom ratings or with 7 of 9 cognitive measures. The relationship between the number of Na^+^ peaks and PANSS Hallucinations is shown in Figure 4b (p=0.0033, coeff.=0.77). In striking contrast, sEPSC amplitude showed no correlation with positive symptoms, but was positively correlated with ratings for multiple negative symptoms, and negatively correlated with Factor 4 Processing Speed, Factor 5 Card Sorting (WCST), and IQ. Figure 4c shows the positive correlation between sEPSC amplitude and PANSS Emotional Withdrawal (p=0.0071, coeff.=0.73), and Figure 4d shows the negative correlation (p=2.94E-05, coeff.=-0.93) between sEPSC amplitude and performance on WCST. Note that these results appear independent of PRS, which is uncorrelated with sEPSC amplitude and weakly *negatively* correlated with the number of Na^+^ peaks (Figure 4a). This lack of correlation is not surprising given the compressed range of variation of PRS in patients. Results for SCZ and CON, including the adjusted p values, for 7 electrophysiological measures and all clinical and cognitive phenotypes tested are shown in **Supplementary Tables 3 and 4**.

**Figure 4.**
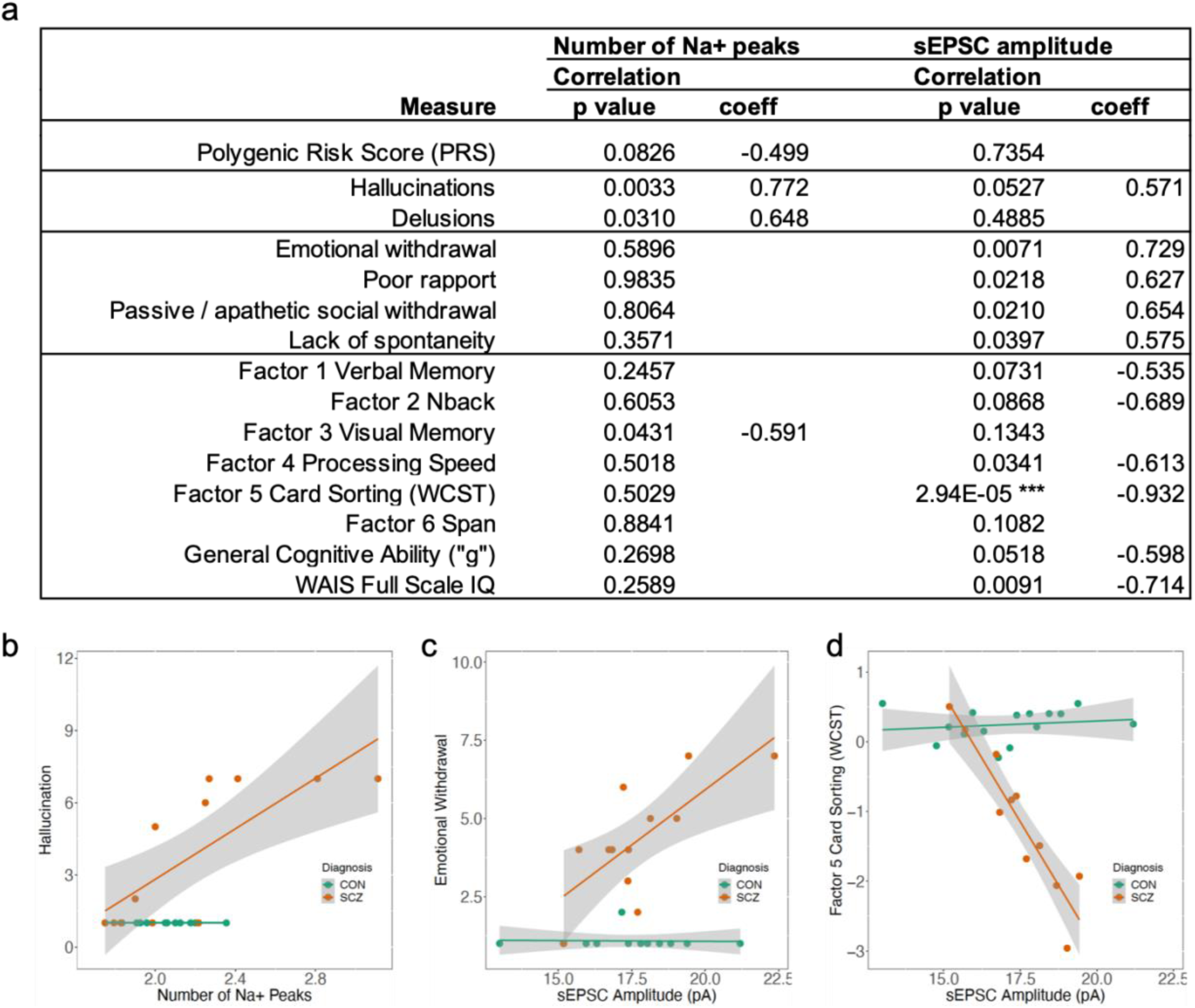
Association between two electrophysiological phenotypes and selected clinical and cognitive phenotypes. **a,** Correlation results in 13 SCZ. Shown are all 8 cognitive phenotypes tested, along with the clinical phenotypes with p<0.05 with either electrophysiological measure. ***adjusted p=0.00014. Results for SCZ and CON, including the adjusted p values, for 7 electrophysiological measures and all clinical and cognitive phenotypes tested are shown in Supplementary Tables 3 and 4. **b,** Relationships between the number of Na^+^ peaks and a positive symptom, hallucination (CON N=12, p not calculated; SCZ N=12, R=0.77, p=0.0033). **c,** Relationships between sEPSC amplitude and a negative symptom, emotional withdrawal (CON N=12, R=-0.034, p=0.92; SCZ N=12, R=0.73, p=0.0071). **d,** Relationships between sEPSC amplitude and Wisconsin Card Sorting Test (WCST) performance (CON N=15, R=0.16, p=0.58; SCZ N=11, R=-0.93, p=2.94E-05).

Linear regression results for CON and SCZ together are shown in **Supplementary Table 5**. Most notable is the association between sEPSC amplitude and WCST in the combined SCZ+CON sample, with a highly significant main effect across groups (p=7.75E-08, adjusted R^2^=0.8947) and interaction of sEPSC amplitude with diagnostic group (p=1.23E-08). Given the small sample sizes, and the facts that subject selection and generation of cellular measures were blind to patient phenotypes, the strength and consistent linearity of some of the observed associations, and their specificity to SCZ, is noteworthy.

## Discussion

hiPSC-derived cellular models have revolutionized the experimental biology of human disease. In recent years, an increasing number of studies of a variety of disorders with pathognomonic cellular pathology have used hiPSCs to elucidate etiologic and pathogenic mechanisms^31–33^. In the case of psychiatric disorders, however, the challenge to model illness at the cellular and circuit level is particularly difficult because there are no well-established pathological cellular characteristics, no evidence of autophagy or apoptosis, no inclusion bodies or even consistent molecular associations. Moreover, no prior study has demonstrated a robust association between an hiPSC-derived neuronal measure and clinically relevant features of the psychiatric disorder being modeled.

In the present report, in an effort to bridge the gap between experimental measures and clinically meaningful characteristics of the individual donors, we have taken a classic translational approach, exploring a diverse landscape of physiological measures in hiPSC-derived neurons from individuals with SCZ who have been deeply characterized clinically. To maximize the biological distance between groups, we used genomic information to select individuals in the SCZ and neurotypical groups such that their PRS distributions did not overlap. We report several illness-associated physiological phenotypes in hiPSC-derived neurons that predict cardinal clinical manifestations of illness. Further, these associations aggregate in patterns that are consistent with clinical experience and well-supported in clinical research.

We observed no significant, reproducible differences between SCZ and CON lines in many important cellular phenotypes including pluripotency, number and type of neurons produced, and network activity as measured by Ca^2+^ imaging. However, multiple electrophysiological measures did show differences by diagnosis, which when taken together suggested that SCZ neurons display altered Na^+^ channel function. These phenotypes included increased membrane resistance, decreased Na^+^ current threshold, increased frequency of Na^+^ currents generated in response to a voltage ramp and hyperpolarized shifts in the activation and inactivation kinetics of Na^+^ currents, collectively indicating Na^+^ channel function is altered in cortical neurons derived from individuals with SCZ compared to CON. These results are in agreement with several previous transcriptomic studies from postmortem human brains, where DEGs were enriched in pathways associated with ion channel function^5,34–36^.

Although these phenotypes implicate Na^+^ channel dysfunction, whether the observed effects reflect differential *α* and *β* subunit expression and/or differential channel modulation via intracellular signalling pathways remains to be determined. In addition to changes in Na^+^ channels, the frequency of GABA transmission (sIPSC) was increased in neurons from SCZ individuals while glutamatergic transmission remained unchanged (sEPSC), suggesting alteration in synaptic excitatory/inhibitory balance. Given that the fraction of GABA-ergic cells, the expression of chloride transporters, and the frequency of network activity did not differ by diagnosis, we hypothesize that this phenotype may reflect an increase in the number of inhibitory GABA-ergic synapses or an alteration in the probability of transmitter release. These findings are consistent with a previous SCZ hiPSC study showing that neuronal DEGs were associated with an upregulation of GABA-ergic synapses^18^, which could contribute to the altered oscillatory activity that has been associated with SCZ^37^.

In addition to the physiological measures, we identified 68 DEGs, many of which were previously identified in other studies using hiPSCs. We observed dysregulation of two protocadherins (*PCDHA5* up, *PCDHA6* down), members of a family of genes previously associated with schizophrenia^38^ and shown to be disrupted in cortical interneurons derived from SCZ patients^16^. Knockout of the entire *Pcdha* locus in mice reduced the number of inhibitory synapses^16^, and hence upregulation of *PCDHA5* in SCZ may contribute to the observed increase in sIPSC frequency. Gene set enrichment analyses suggested involvement of pathways related to WNT signaling and forebrain development. Dysregulation of both of these pathways has been consistently observed in previous studies of SCZ using hiPSCs^16, 39–41^. In addition, we found that NPC DEGs correlated with neuronal DEGs, which is also consistent with previous findings^10^. Together, these findings provide further support for an early neurodevelopmental component to SCZ^42^, and for WNT signaling and protocadherin biology as candidate pathways contributing to SCZ etiology.

A central advantage of this study was the unique use of legacy data from our study where we conducted extensive and rigorous clinical characterization and tests of cognitive performance^25, 27^. The exploratory study design reported here allowed us to test associations of cellular phenotypes in patient cell lines with the behavioral and cognitive phenotypes of the actual donors. Due to practical limitations, the sample sizes were small and because there were no similar prior studies, we could not reasonably estimate expected effect sizes or error rates. Based on the correlation structure of the variables, we adjusted the p values for the effective number of tests performed. After this adjustment (and similarly, after Bonferroni correction), only three associations remained with p<0.05: sIPSC frequency with diagnosis (adjusted p=3.86E-05; **Supplementary Table 2**), Na^+^ inactivation with diagnosis (adjusted p<0.0035; Fig. 3f), and sEPSC amplitude with Wisconsin Card Sort Test performance in SCZ (adjusted p=0.0001; **Supplementary Table 3**), with the latter showing a remarkably clear, almost monotonic relationship. Nevertheless, it is the broader *pattern* of association among cognitive, positive, and negative symptom variables (including some that did not “survive correction”) that is noteworthy and merits further investigation. Importantly, the secondary analyses and the clustering of phenotypes pointing to Na^+^ channel kinetics strengthens our interpretation of the findings.

The unexpectedly strong associations from these small sample sizes supports the notion that the availability of deeply clinically phenotyped donor cohorts is critical for testing the potential validity of electrophysiological phenotypes of patient-derived neurons. Not only were the effects relatively strong but the patterns of association were divergent and aligned with clinical literature in interesting ways. The number of Na^+^ peaks showed associations with two (correlated) canonical positive symptoms of SCZ, hallucinations and delusions, but no association with negative symptomatology and little with cognitive performance. In contrast, sEPSC amplitude showed a broad association with negative symptoms and cognitive dysfunction, but no connection to positive symptoms. This divergence recalls a substantial body of work implicating relatively consistent clinical subgroupings of individuals with SCZ (including positive vs negative/cognitive) and distinct biological underpinnings for these subgroups^43^.

The novel study design also included elements designed to monitor and reduce unwanted variability and artifacts. By comparing high PRS SCZ and low PRS CON, we aimed to magnify cellular and network phenotypes that are driven by genomic risk for SCZ, since these phenotypes are specifically what we seek to disentangle at the mechanistic level. Subjects were males of European ancestry, chosen on PRS and general availability of genomic and phenotypic data, but there were no selection criteria applied to stratify by specific clinical or cognitive features. All experiments were conducted blind to diagnosis and cell lines were handled in random SCZ-CON pairs. To minimize batch effects, we conducted at least 4 rounds of differentiation, with 2 experimenters in rotation. These combined elements were designed to reduce experimental bias, and thus increase confidence in the measurements obtained. Nevertheless, due to the small sample sizes and the number of tests performed, some of the associations observed may be spurious, so replication studies with larger samples are necessary to confirm and extend these findings.

In summary, these data provide the first putative associations in SCZ between specific electrophysiological measures in human neurons and cardinal clinical and cognitive phenotypes of the adult donors from whom the cells originated. Integration of insights from a wider range of data types may provide the synergy necessary for elucidation of biological risk mechanisms, as well as the identification of pathological molecular and cellular pathways and biomarkers as leads towards personalized medicine.

## Methods

### Subjects and fibroblasts sampling

Subjects were males of European ancestry from the Sibling Study of Schizophrenia at the National Institute of Mental Health in the Clinical Brain Disorders Branch (NIMH, protocol 95M0150, NCT00001486, Annual Report number: ZIA MH002942053, DRW PI) with additional support from the Clinical Translational Neuroscience Branch, NIMH (KFB PI)^25^, chosen solely on their PRS scores and the availability of clinical and cognitive data^27^. All subjects were extensively screened by obtaining medical, psychiatric and neurological histories, physical examinations, MRI scans, and genome-wide genotyping to rule out diagnosable clinical disorders. Following informed consent, skin fibroblasts were cultured from biopsies taken from the mesial aspect of the upper arm using a 3mm diameter punch under lidocaine intradermal anesthesia.

### Polygenic Risk Score (PRS) Calculation

Polygenic Risk Scores (PRS) are a sample dependent estimate of an individual’s genomic risk for SCZ. PRS were calculated based on the most commonly used “PGC2” Psychiatric Genomics Consortium GWAS for SCZ^44^. We obtained odds ratios of 102,217 index SNPs from a meta-analysis of PGC2 GWAS using datasets excluding our contributed sample. These 102,217 SNPs are LD independent (R^2^<0.1) and span the genome. We then calculated PRS, a weighted sum of risk alleles, by summing the imputation probability for the reference allele of the index SNP, weighted by the natural log of the odds ratio of association, as described^44^. Ten PRS (PRS1-PRS10) were calculated using SNP subsets under different thresholds of the PGC2 GWAS p-values of association with SCZ: 5E-08, 1E-06, 1E-04, 0.001, 0.01, 0.05, 0.1, 0.2, 0.5, and 1. SNPs in sets with lesser p-values are contained in sets with greater p-values (SNPs in PRS1 are included in PRS2, SNPs in PRS1 and PRS2 are included in PRS3, etc.). We chose fibroblast lines to reprogram based on PRS6, which was constructed using 24,670 SNPs with GWAS association p values less than 0.05, since this threshold had the greatest prediction accuracy for diagnostic status in multiple independent samples^44^.

### Reprogramming of fibroblasts into hiPSCs

Early passage fibroblasts (less than passage 5) were thawed and cultured in cDMEM media (DMEM, 10% Fetal Bovine Serum, 1% Non-Essential Amino Acids (NEAA), beta-mercaptoethanol (b-ME) and 1% Penicillin/Streptomycin). According to the manufacturer’s protocol ^45^, all cell lines were reprogrammed using Cytotune^TM^-iPS 2.0 Sendai Reprogramming Kit (A16517; ThermoFisher Scientific). Clones obtained were cultured onto irradiated mouse embryonic fibroblast (ir-MEF) feeder layer in hESC media (DMEM/F-12, 20% Knockout serum replacement (KSR), 1% NEAA, b-ME) supplemented with FGF2 (10ng/ml; 100-18B; Peprotech) with daily medium changes. iPS cells were passaged once a week (∼85-90% confluent wells) using collagenase, typeIV (1mg/ml; 17104019; Gibco).

### Validation of hiPSC pluripotency

For stem cell quality control, immunocytochemistry of pluripotency markers (NANOG, SOX2, OCT3/4 and TRA-1-60), quantitative PCR for pluripotency markers (NANOG, OCT3/4, SOX2, KLF4 and LIN28), Karyotyping (G-banding) and Alkaline Phosphatase (AP) staining (data not shown) was performed on all iPS cell lines used in the study. For pluripotency marker staining, iPS cells were fixed with 4% paraformaldehyde for 15 minutes. Permeabilization and blocking was performed simultaneously with 0.3% Triton X-100 and 10% donkey serum (blocking buffer) by incubating for at least 45 minutes at room temperature. Primary antibodies against NANOG, SOX2, OCT3/4 and TRA-1-60 were diluted in the blocking buffer and incubated overnight at 4° C. After three 5-minute DPBS washes, cells were incubated with corresponding fluorescent-conjugated secondary antibodies for 2 hours at room temperature. Three 5-minute DPBS washes were performed and cells were counterstained with DAPI (4′,6-diamidino-2-phenylindole). All imaging was performed at 10X on Operetta (see below). For quantitative PCR, total RNA was extracted from iPS cells using phenol:chloroform isolation with Trizol followed by purification using the RNeasy Micro Prep kit (Qiagen). Subsequently, cDNA was prepared using High-Capacity cDNA Reverse Transcription Kit (4387406; Thermofisher Scientific). For karyotyping analysis, cells were grown on 25cm^2^ tissue culture flasks and karyotyping was performed at WiCell (chromosome analysis service). For AP staining, cells were plated on 12-well plates. Live AP staining (A14353; Thermofisher Scientific) was performed and AP activity was imaged using standard FITC filter.

### Differentiation into cortical neurons

iPS cells were differentiated into cortical neurons as previously described in^12^ with modifications. Briefly, iPS colonies were detached from the ir-MEF feeder layer by incubating the cells with collagenase, type IV (1mg/ml; Gibco) for 1h at 37° C. The detached colonies were cultured in suspension in ultra-low attachment 10 cm culture dishes (Corning) in a FGF-2 free hESC media supplemented with Dorsomorphin (2uM) and A-83 (2uM) (here on referred to as embryoid body (EB) media) for 4 days with daily medium changes. On day 5, EB media was replaced with neural induction (hNPC) media consisting of DMEM/F-12, N2 supplement and NEAA supplemented with heparin (2mg/ml) and cyclopamine (2uM). On day 7, EBs growing in suspension were then transferred onto Matrigel-coated 10 cm dishes to form neural tube-like rosettes. The attached rosettes were cultured for 10 days with media changes every other day. After 10 days, rosettes were mechanically picked using hand-pulled glass pasteur pipettes and transferred to ultra-low attachment 6 well plates. Picked rosettes were cultured for additional 3 days in hNPC medium containing B27 and Penicillin/Streptomycin forming neural spheres. For neuronal differentiation, neural spheres were dissociated with Accutase (Gibco) at 37° C for 10 min. Cells were passed through a 40um mesh filter to obtain a single cell suspension of neural progenitor cells (NPCs). NPCs were then plated onto plates containing a confluent layer of rat primary astrocytes (see below). Neuronal cultures were treated with cytosine arabinoside treatment (AraC; 4uM) after 1 week in culture (DIV7) for 7 days to DIV14. Neuronal cultures were maintained for up to 10 weeks (DIV70) with partial (½) medium changes twice a week with Neuron medium (Neurobasal media, 1% Glutamax, B27 supplement supplemented with Ascorbic Acid (200nM), cAMP (1uM), BDNF (10mg/ml) and GDNF (10mg/ml)). Neurons were plated on 24-well or 96-well ibidi plates for subsequent live imaging or high-content imaging, on glass coverslips for electrophysiology or live imaging, or standard 24 well plates for biochemical assessments.

### Primary rat astrocytes

Primary astrocytes were prepared as previously described with modifications^46, 47^. Cortices from a litter of 2-3 day old pups were isolated. After dissection of the tissue and removal of the meninges, the cortices were triturated with a 1ml pipette 10 times to break down the tissue into smaller pieces. The tissue was then incubated with 300µl 2.5% trypsin-EDTA (Gibco) and 100µl DNase (100ug/ml; Sigma) on a shaker at 150 rpm at 37° C for 30-45 minutes. Subsequently, the cells were passed through a 45µm mesh filter, and transferred in Glial media (MEM, 5% FBS, Glucose (20mM), Sodium bicarbonate (0.2mg/ml), L-Glutamine (2.5mM) and 1% Penicillin/Streptomycin) to 75cm^2^ tissue culture flasks (5-6 cortices per flask). After 24h, the flasks were vigorously tapped and shaken to remove any loosely attached cells. The cells were cultured for 6-7 days with medium changes every other day. After 7 days, flasks were 80-85% confluent and were passaged onto poly-D-lysine/Laminin coated 24-well (seeding density 70,000 cells/well) or 96-well plates (seeding density 20,000 cells/well). Plated astrocytes were mitotically arrested via AraC (20uM) 24h after plating, prior to adding human NPCs.

### Immunocytochemistry

For neuronal staining, samples were fixed with ice-cold 4% paraformaldehyde for 10 min. Permeabilization and blocking were performed simultaneously with 3-5% normal goat serum, and antigen-retrieval, if indicated, was performed by a 60 minute incubation in sodium citrate buffer at 80° C. Primary antibodies directed against NeuN, CTIP2, TBR1, FOXP2, SATB2, CUX1, BRN2, GAD67, TH were prepared in the blocking buffer and incubated with the sample at 4° C overnight. Fluorescently conjugated secondary antibodies were prepared in blocking buffer and incubated with the sample at room temperature for 90 minutes. Samples were counterstained with DAPI and stored in PBS until imaging.

### High content imaging and analysis

For neuronal identity markers, at least 25 randomly located images per well were acquired with a 20X/0.6NA objective on an Operetta (PerkinElmer) and processed using custom code in Columbus (PerkinElmer). DAPI segmentation was used to identify cell bodies, and NeuN segmentation was used to identify regions of interest (ROIs) as neurons. Neuronal subtype markers (BRN2, CTIP2, CUX1, FOXP2, GAD67, SATB2, TBR1, TH) were identified based on colocalization with DAPI+NEUN+ ROIs. Analysis was performed on final cell counts in R using full linear models (for neuron identity markers) or linear mixed effects models (for NeuN).

### RNA isolation and qPCR

Total RNA was isolated as described above, quantified using a Nanodrop spectrophotometer, and 100ng was reverse-transcribed using Superscript III. For neuron quality control, quantitative PCR was performed using the Fluidigm Biomark system with Taqman probes according to manufacturer’s instructions. We utilized human specific Taqman probes that would not bind to rat sequence to validate gene expression in the human transcriptome, and rat specific Taqman probes that would not bind to the human sequence to validated gene expression in the rat transcriptome, and verified that these probes performed as expected by using controls from each species independently. Briefly, cDNA was amplified for 14 cycles of PCR using a combination of all included probes, then the preamplified product was assayed in duplicate for each of 90 genes of interest, three housekeeping probes, and negative controls. The mean of the Ct values for the two technical replicates was computed and normalized against the mean of the Ct values for all three housekeeping genes.

### RNA sequencing data generation and processing

Illumina RNA sequencing (RNA-seq) libraries were constructed with Ribo-Zero gold kits across three batches and included ERCC spike-in sequences. The first batch-which only included hiPSC and NPC samples - had libraries prepared and sequenced at the Sequencing Core at the Lieber Institute for Brain Development on an Illumina HiSeq 3000. The second and third batches - which included all neuronal samples and 5 repeated NPC samples for comparability - were prepared and sequenced at Psomagen (formerly Macrogen) on an Illumina HiSeq 4000. A total of 304 samples were sequenced with paired end 100bp reads (2×100). Raw sequencing reads from all samples were aligned to a custom concatenated Gencode hg38 + rn6 reference genome using HISAT2 2.0.4^48^ (including NPC samples without any rodent RNA, for comparability to neuronal samples). Genes and exons were quantified with featureCounts v1.5.0-p3^49^ to a custom concatenated hg38+rn6 gtf file within each sample. ERCC spike-in sequences were quantified with kallisto^50^, and we further calculated a bias factor for each sample using the observed versus expected abundances [∑((obs-exp)^2)]. A barcode of 738 exonic/coding SNPs were genotyped using the RNA sequencing data to confirm sample identities using samtools mpileup^51^. We used a previous RNA deconvolution method developed by our group to estimate the RNA fractions for 10 different cell classes in each sample from gene expression levels^9^.

After processing, we performed quality control checks of samples for RNA quality, sequencing quality, and sample identities. We examined the distributions of read alignment and gene assignment rates to both human and rodent genomes, as well as the ERCC spike-in accuracy rates (sum of squared deviations from expected concentrations). We confirmed the sample identities within and across subclonal lines using the RNA-derived genotype barcodes (for donor confirmation) and RNA fractions (for timepoint confirmation).

### RNA-seq data analysis

After quality control checks, we performed a series of analyses on different subsets of samples.

1. All samples (N=304) for time-course/developmental changes
2. Neuronal samples (N=94) for case-control modeling, including 51 CON samples (from 18 subclonal lines from 14 genomes) and 43 SCZ samples (from 14 lines from 12 genomes) across 23,119 expressed genes (based on subset-specific RPKM>0.2)
3. NPC samples (N=143) for case control modeling, including 75 CON (from 17 subclones from 13 genomes) versus 68 SCZ samples (from 16 subclones from 14 genomes) across 23,974 expressed genes (based on subset-specific RPKM>0.2)

Principal component analysis (PCA) of normalized gene expression levels (log2[RPKM+1]) was used for exploratory data analysis within each subset of samples. The top component in each subset of samples (PC1) related to underlying RNA composition (which reflected developmental changes in the full dataset) and we therefore constructed the top principal component of these 10 RNA fractions to capture their correlated effects (“cellPC”, see Extended Data Figure 4). We used linear mixed effects modeling, treating each subclone line as a random intercept, using the limma voom approach^20^. In neuronal samples, we specifically modeled TMM-normalized expression as a function of diagnosis, adjusting for main effects of ERCC bias (quantitative), cellPC (quantitative), DIV (binary, 8 or 10 weeks), the human alignment fraction (quantitative), and the human gene assignment rate, and random intercept of subclone line. In NPC samples, we used a related statistical model for diagnosis that adjusted for DIV, cellPC, chrM mapping rate (quantitative), human gene assignment rate, ERCC bias, and sequencing batch, with subclone again modeled as a random intercept. For both analyses, we calculated log2 fold changes for the SCZ term (relative to CON, such that positive log fold changes indicated increased expression in patients), which were used to calculate empirical Bayes t-statistics and corresponding p-values [limma]. P-values were adjusted for multiple testing using the Benjamini-Hochberg (B-H) procedure to control for a false discovery rate (FDR) of 0.1. Gene set enrichment analyses were performed with the clusterProfiler Bioconductor package^52^, using the hypergeometric test to assess enrichment of marginally-significant differentially expressed genes (at p<0.005) that were stratified by case-control directionality, against a background of all expressed genes that had Entrez gene IDs. We again used the B-H FDR to control for multiple testing of gene sets.

### Analysis of association between electrophysiological measures and diagnosis

We performed linear regression analyses between 47 electrophysiology measures and schizophrenia diagnosis. The complete results are shown in **Supplementary Table 2**. We adjusted for the researcher who differentiated the neurons (two people/levels/groups), the researcher who recorded from the neurons (three people/levels/groups), the developmental time-point of the neurons (binarized into early or late, two groups). We opted not to adjust for the donor/cell line of each neuron due to the design matrix being singular/collinear, which resulted in directional instabilities of the schizophrenia effect (e.g. different factor orderings of the Line term resulted in different directionalities of the schizophrenia effect on the ePhys measurements). Secondary analyses utilized Tukey post-hoc tests to ensure directional/model consistency, as well as timepoint-specific sensitivity analyses to ensure results were not driven by a single time point. As many of the 47 variables were correlated, to calculate p values adjusted for the effective number of tests, we applied the Meff correction factor^53, 54^, which was 34.91.

### Analysis of association between electrophysiological measures and clinical and cognitive data

In order to produce a single (genome-based) value for each subject for the 7 electrophysiological phenotypes, we took the mean values across both 8 (DIV56) and 10 week (DIV70) time points, and across lines. Pearson correlations tested linear associations with clinical and cognitive variables separately for 13 SCZ and 15 CON. The complete results are shown in **Supplementary Tables 3** (SCZ) and **4** (CON). These 7 electrophysiological variables, themselves, were largely uncorrelated. The clinical (PANSS) and cognitive variables are each correlated, so we applied the Meff correction: for the 30 PANSS phenotypes, the correction factor was 24.24, and for the 8 cognitive variables, the correction factor was 4.88. The linear regression p values shown in **Supplementary Table 5** are uncorrected.

### Virus infection and Calcium Imaging and analysis

Neurons were infected at DIV23 with AAV1-hSyn1-mRuby2-P2A-GCaMP6s (a gift from Tobias Bonhoeffer, Mark Huebener and Tobias Rose (Addgene viral prep # 50942-AAV1; http://n2t.net/addgene:50942; RRID:Addgene_50942)^55^. Imaging was performed in culture media at DIV42 and DIV63 on a Zeiss LSM780 equipped with a 10X/0.45NA objective and a temperature- and atmospheric-controlled enclosure. A reference image was acquired for each field of mRuby fluorescence, then a time-series was acquired at 4Hz for 8 minutes. Regions of interest were identified from the red reference image, and fluorescent intensity over time was computed for each ROI then normalized to ΔF/F. Peaks were identified and the number of events was calculated for each ROI. The synchronicity of these events was correlated on a per-image basis. In some cases, pharmacological inhibitors were then added to block glutamatergic synaptic transmission by application of dl-AP5 (100uM) and DNQX (10uM).

### Whole-cell patch-clamp recordings

Whole-cell electrophysiology recordings were performed using the following solutions. Extracellular recording buffer contained (in mM): 128 NaCl, 30 glucose, 25 HEPES, 5 KCl, 2 CaCl_2_, and 1 MgCl_2_ (pH 7.3). Patch pipettes were fabricated from borosilicate glass (N51A, King Precision Glass, Inc.) to a resistance of 2-5 MΩ. For the majority of current-and voltage-clamp measurements, pipettes were filled with (in mM): 135 KGluconate, 10 Tris-phosphocreatine, 10 HEPES, 5 EGTA, 4 MgATP, and 0.5 Na_2_GTP (pH 7.3). For Na^+^ channel kinetics experiments (Fig. 3) pipettes were filled with (in mM) 10 NaCl, 140 CsF, 10 HEPES, 1 EGTA, 15 D-Glucose, 10 TEA-Cl (ph 7.3). Extracellular recording buffer (in mM) 40 NaCl, 75 Choline-Cl, 1 CaCl_2_, 1 MgCl2, 10 HEPES, 20 TEA-Cl, 4AP, 0.1 CdCl_2_, 10 D-Glucose (pH 7.4). Current signals were recorded with either an Axopatch 200B (Molecular Devices) or a Multiclamp 700A amplifier (Molecular Devices) and were filtered at 2 kHz using a built in Bessel filter and digitized at 10 kHz. Voltage signals were filtered at 2kHz and digitized at 10kHz. Data was acquired using Axograph on a Dell PC (Windows 7). For voltage clamp recordings, cells were held at −70 mV for sodium/potassium currents and sEPSCs, and at 0 mV for sIPSCs. After recording a ramp protocol (−110 mV to 110 mV, 0.22 mV/s) for each neuron in voltage clamp, the recording configuration was switched to current clamp and a series of action potentials (20 episodes) was elicited at 0.5 Hz with a square current injection that was increased by 10pA every episode. AP onset was defined when the slope exceeded 10 mV/ms. AP threshold and onset time were subsequently identified by Axograph. Baseline was set at onset time (i.e. threshold) to calculate action potential peak amplitude. Width was calculated at 50% of peak amplitude. Calculation of dV/dt max. and min. was set as the maximum slope for the ascending and descending phases with 3 regression points per max slope.

---

**Extended Data Figure 1.**
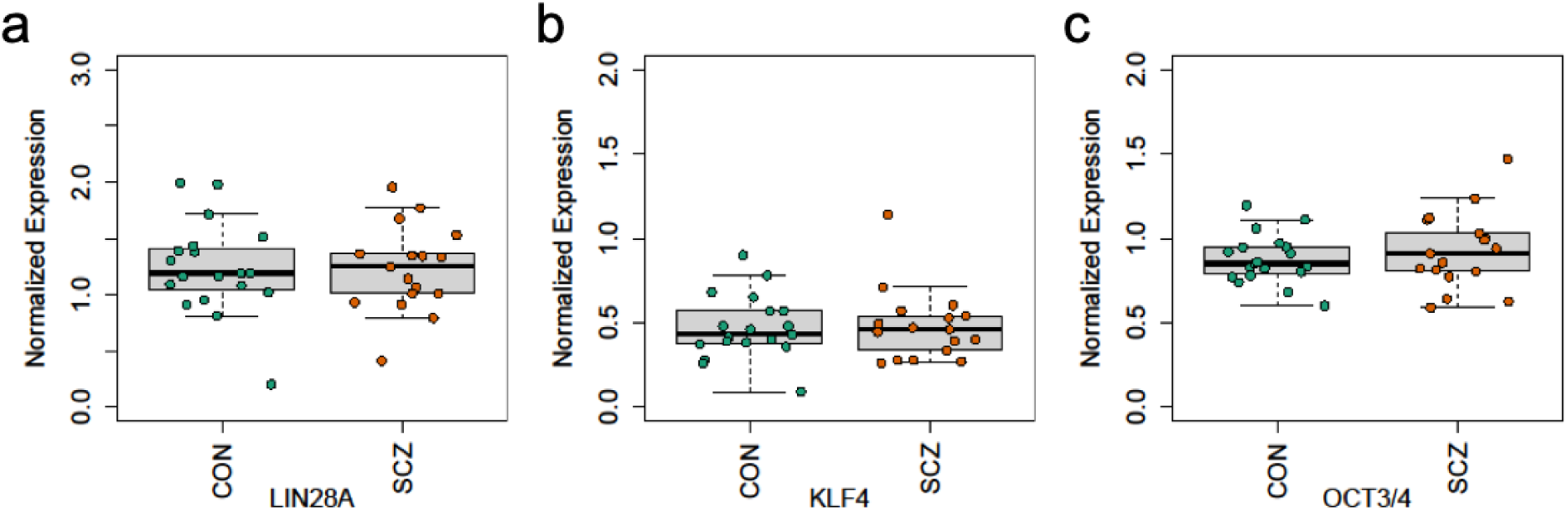
No significant effect of diagnosis on hiPSC pluripotency. qPCR quantification of stem cell markers in **a,** *LIN28A* (CON=1.247 ± 0.101, SCZ=1.225 ± 0.092, p=0.87, N=35 from 28 genomes), **b,** *KLF4* (CON=0.418 ± 0.045, SCZ=0.4805 ± 0.051, p=0.37, N=35 from 28 genomes) and **c,** *OCT3/4* (CON=0.882 ± 0.033, SCZ=0.924 ± 0.055, p=0.51, N=35 from 28 genomes), all normalized to their expression in embryonic stem cell line H1.

**Extended Data Figure 2.**
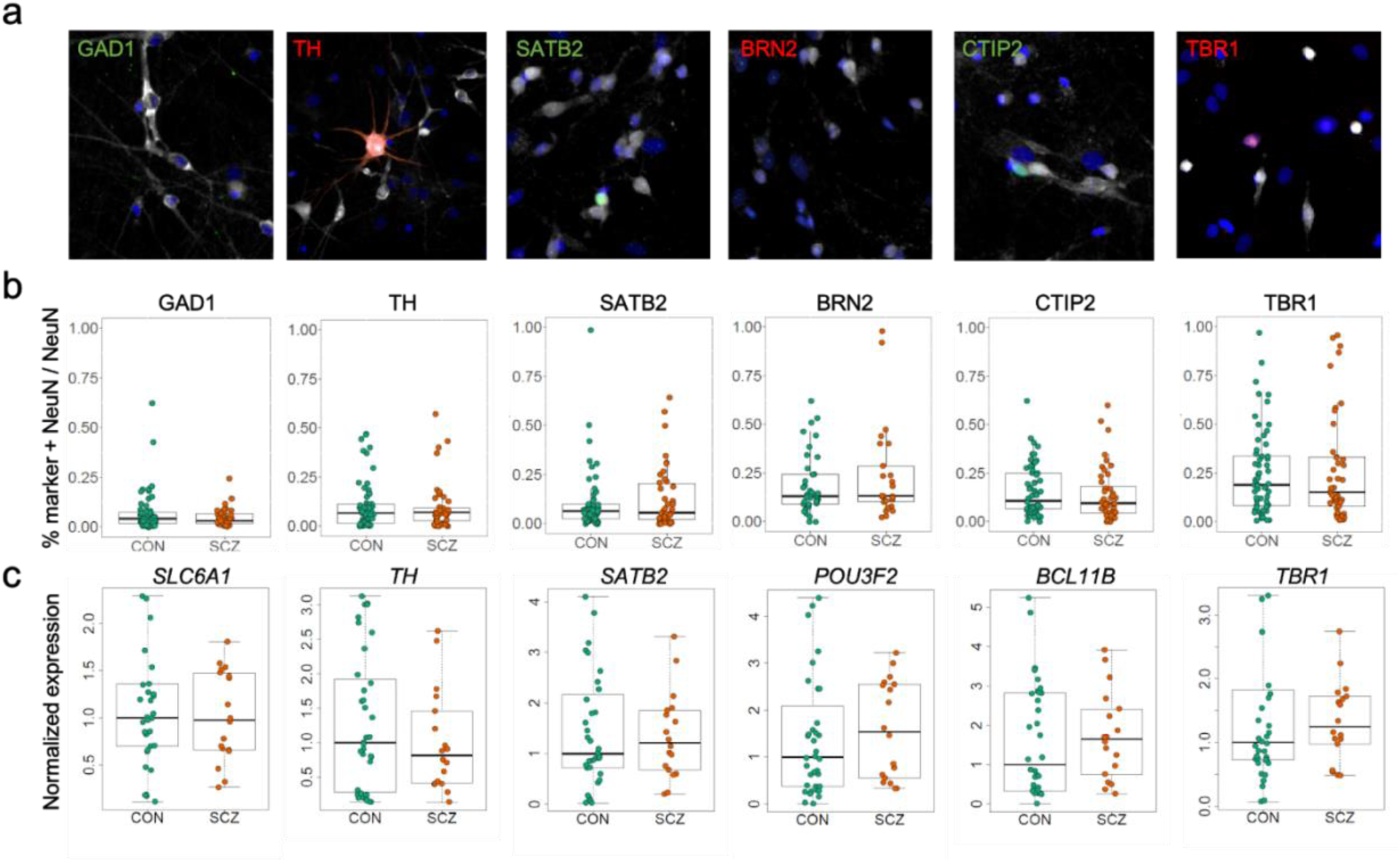
Neuron cell types generated are not different by diagnosis. **a,** Representative image from the same line (orchid), 70 days post-differentiation, DAPI (blue), NeuN (white), and cellular identity markers GAD67 (green) and tyrosine hydroxylase (TH, red), upper cortical layer markers SATB2 (green) and BRN2 (red), and deeper cortical layers markers CTIP2 (green) and TBR1 (red). **b,** Following each differentiation, lines from each cohort were assayed for neuronal identity markers using immunocytochemistry followed by high-content imaging. We observed no differences in cell type composition between SCZ and CON lines (Proportion of marker+NeuN+/NeuN+ cells: GAD1 CON=0.0643 ± 0.015, N=26; SCZ=0.0561 ± 0.013, N=32, full linear model p=0.95; TH CON=0.089 ± 0.018, N=26; SCZ=0.091 ± 0.019, N=29, full linear model p=0.92; SATB2 CON=0.141 ± 0.039, N=27; SCZ=0.107 ± 0.021, N=33, full linear model p=0.003; BRN2 CON=0.192 ± 0.035, N=21; SCZ=0.196 ± 0.040, N=25, full linear model p=0.83; CTIP2 CON=0.182 ± 0.025, N=28; SCZ=0.135 ± 0.021, N=34, full linear model p=0.49; TBR1 CON=0.237 ± 0.042, N=28; SCZ=0.315 ± 0.048, N=34, full linear model p=0.43). **c,** Normalized gene expression at DIV 70 showed no significant differences between CON and SCZ lines. Gene expression was normalized to mean expression of CON: *SLC6A1* (GAT1)(SCZ effect=0.98, p=0.94 N=26 across 11 genomes), *TH* (SCZ effect=0.89, p=0.78 N=27 across 11 genomes, *SATB2* (SCZ effect=1.15, p=0.76, N=27 across 11 genomes), *POU3F2* (BRN2) (SCZ effect=1.49, p=0.43, N=27 across 11 genomes), *BCL11B* (CTIP2) (SCZ effect=1.60, p=0.39, N=27 across 11 genomes), *TBR1* (SCZ effect=1.11, p= 0.78, N=27 across 11 genomes).

**Extended Data Fig. 3.**
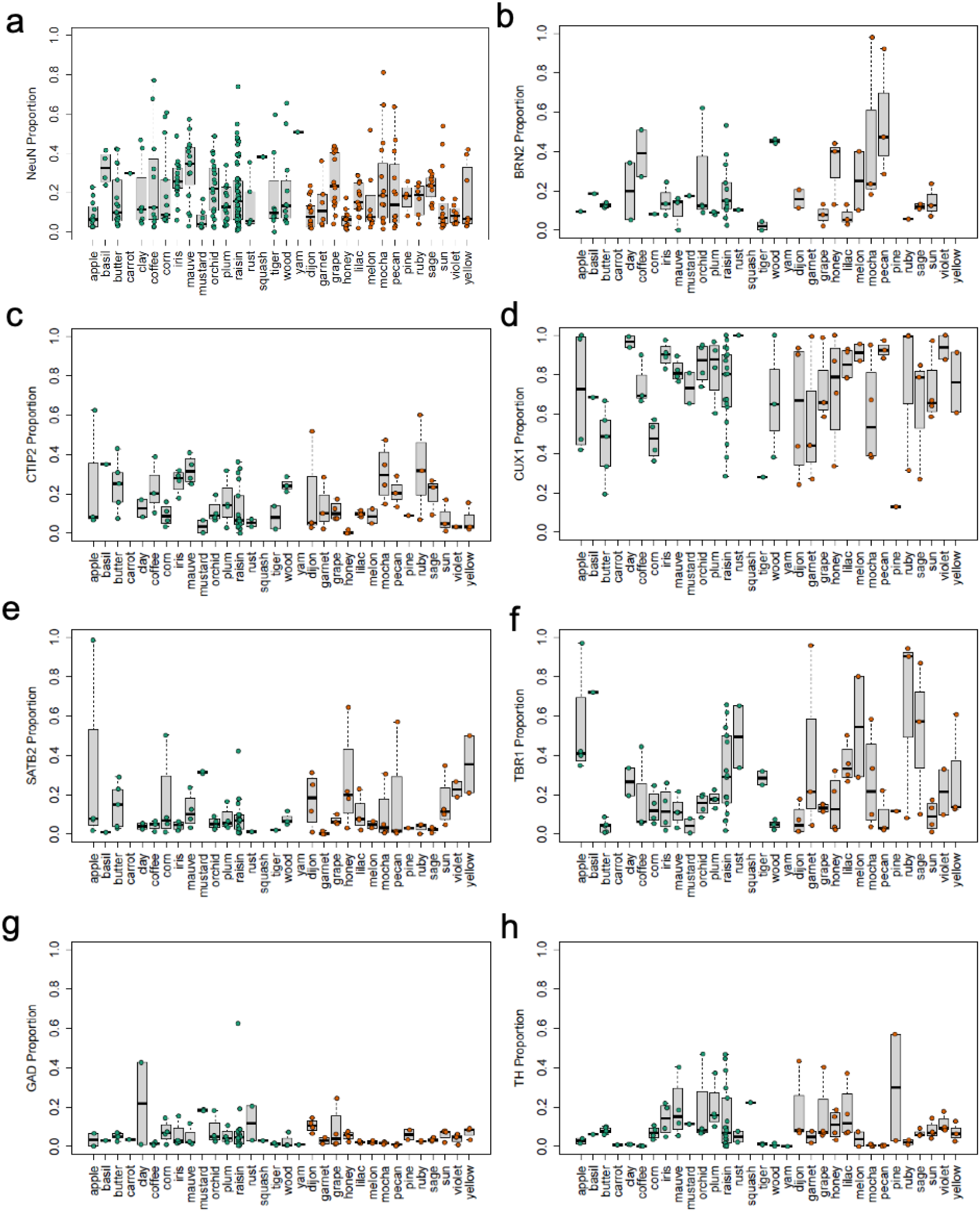
Summary data of immunocytochemistry of neuronal cell type and cortical layer markers for each line. **a,** Summary data by line for percentage of neurons by line. **b,** Summary data by line for percentage of BRN2+ neurons. **c,** Summary data by line for percentage of CTIP2+ neurons. **d,** Summary data by line for percentage of CUX1+ neurons. **e,** Summary data by line for percentage of GABAergic neurons. **f,** Summary data by line for percentage of SATB2+ neurons. **g,** Summary data by line for percentage of TBR1+ neurons. **h,** Summary data by line for percentage of TH+ neurons.

**Extended Data Figure 4.**
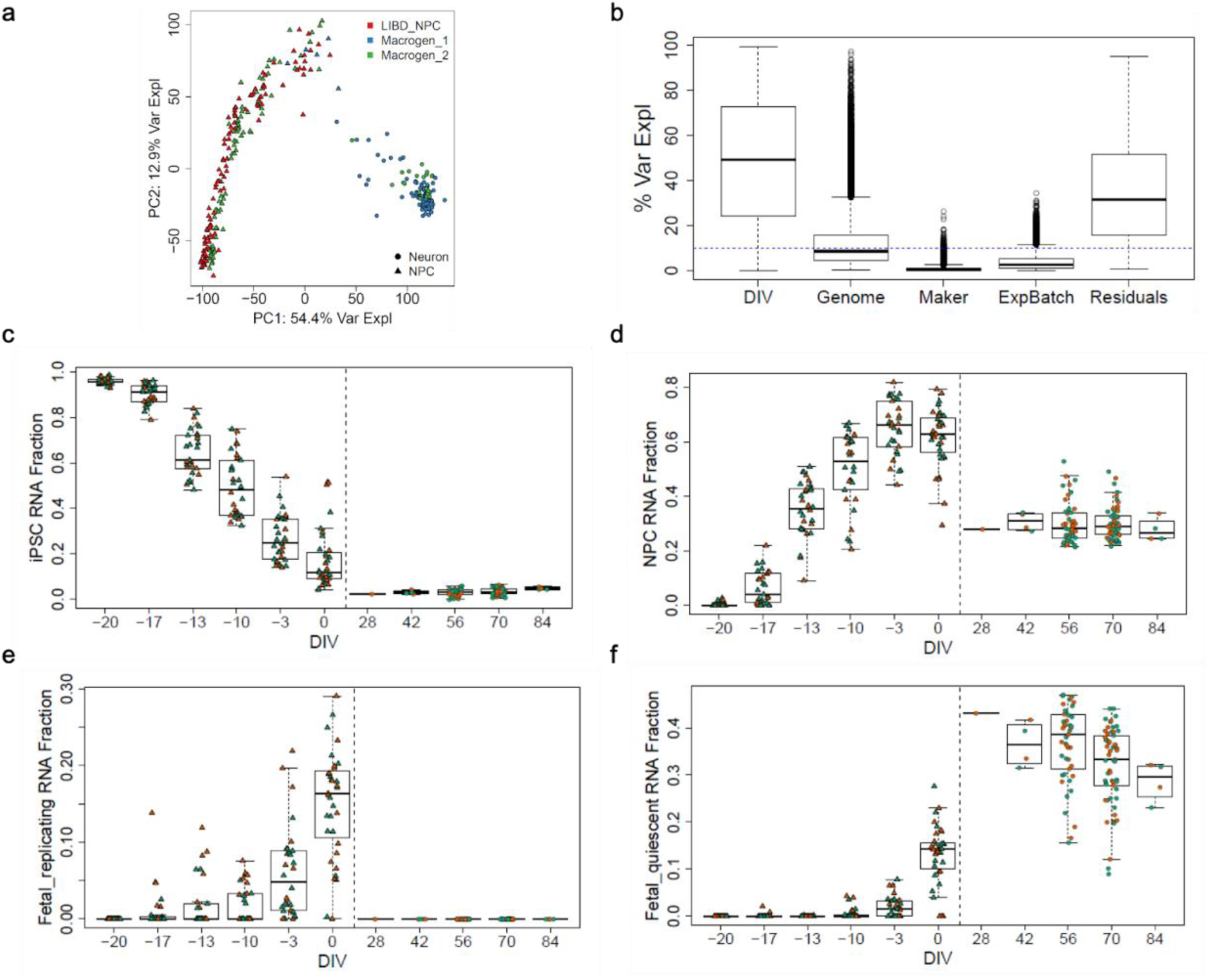
RNA-seq validation of cell states. **a,** Top principal components of gene expression relate to developmental changes. Triangle = NPC, Circle = Neuron; Color = batch. **b,** Variance components analyses of gene expression across each gene (ie each box is the distribution of variance explained across all expressed genes) - developmental stage and genome strongly influence expression variation, while differentiation round or the individual performing the differentiation did not. **c,** We represented cellular heterogeneity as the first principal component of the 10 cell class fractions, which captured 84% of the cellular variance explained, where adult and fetal quiescent neuron RNA fractions were associated with increased maturity, and NPCs and endothelial RNA fractions were associated with decreased maturity. As expected, hiPSC RNA signature decreases over development. **d,** The RNA fraction associated with NPCs increases then decreases over development. **e,** We used RNA deconvolution to estimate the RNA fractions for multiple cell classes, and confirmed that our cells align with the fetal-replicating class during the hNPC stage. **f,** Neurons display the fetal-quiescent class during the post-mitotic neuronal stage.

**Extended Data Figure 5.**
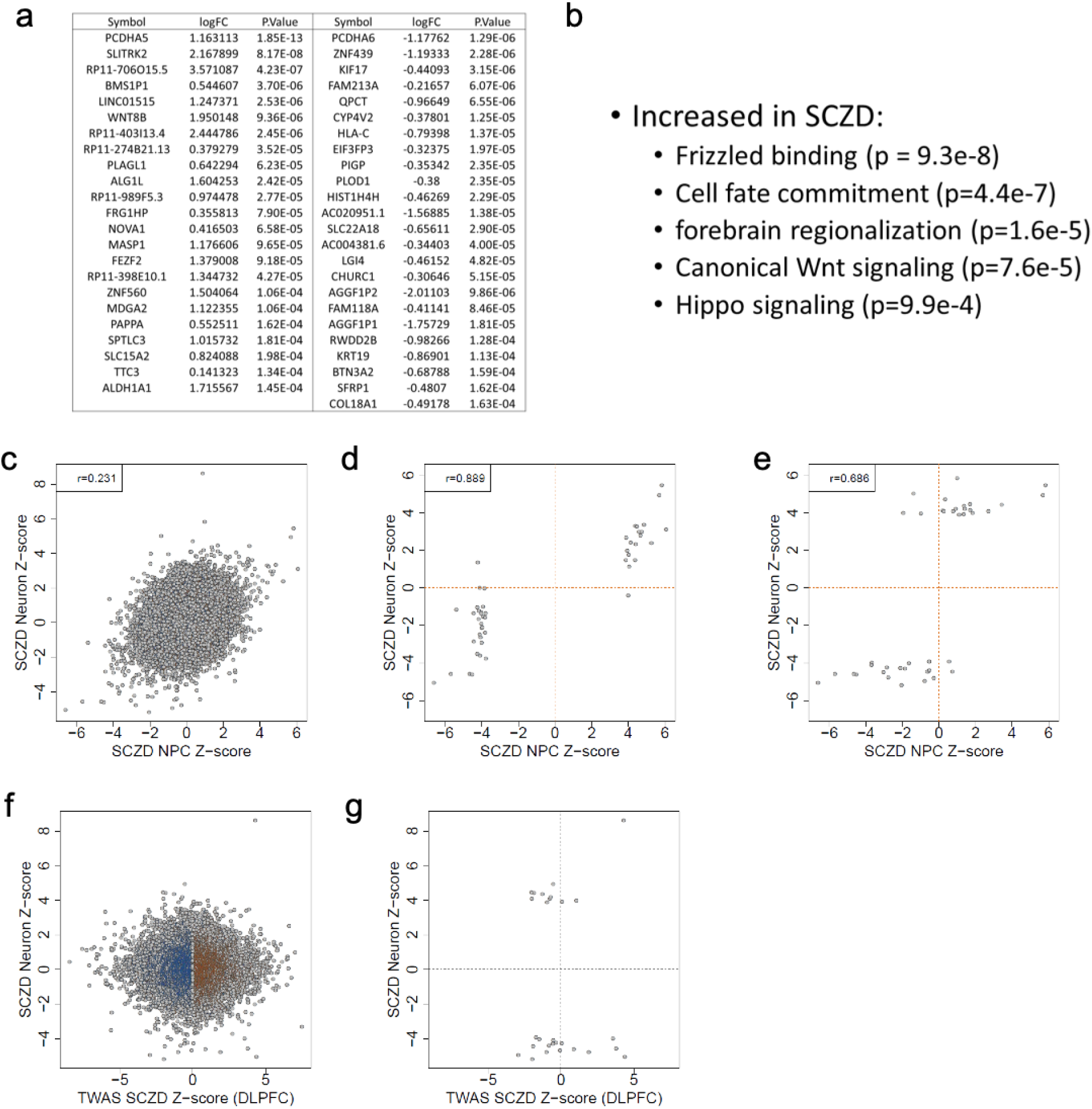
Gene expression changes in hiPSC-derived neurons correlate across cell states. **a,** Differentially expressed genes with FDR<0.05. **b,** Representative GO terms from DEGs at p<0.005. **c,** Gene expression correlates across cell states. **d,** Differentially expressed genes in neurons are observed in NPCs. **e,** Genes that are differentially expressed in NPCs remain differentially expressed in neurons. **f,** Correlation between gene expression in hiPSC-derived neurons when compared to TWAS data from postmortem DLPFC. **g,** Differential expression in hiPSC-derived neurons compared to TWAS data.

**Extended Data Figure 6.**
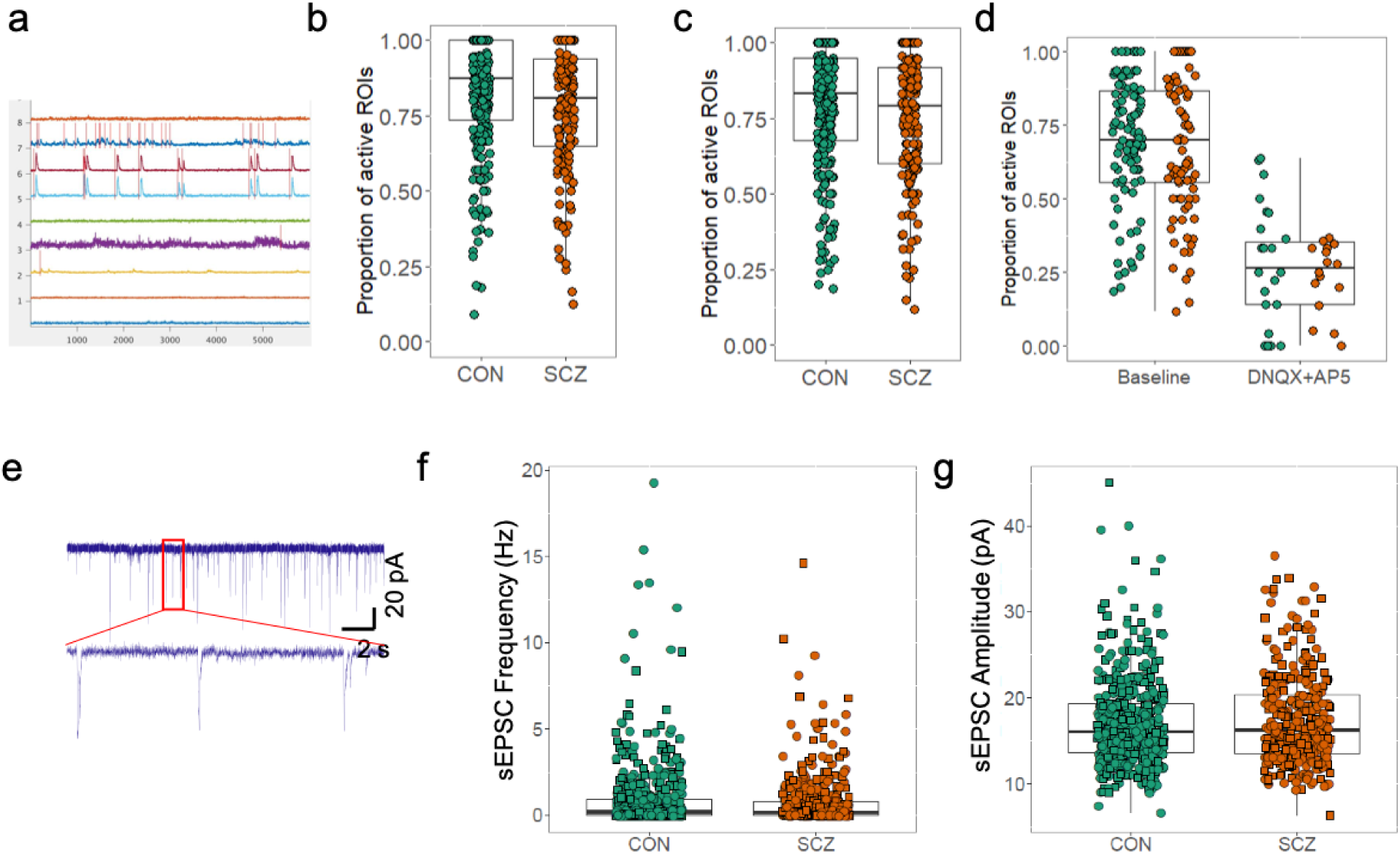
Physiological assessment of cortical neurons. **a,** Representative traces of regions of interest in a control culture. **b,** Proportion of active ROIs are not significantly different between CON and SCZ at an early developmental stage (DIV42, SCZ effect= −0.0494, p=0.403, N=426 across 35 lines). **c,** Proportion of active ROIs are not significantly different at a later developmental stage (DIV63) between CON and SCZ(SCZ effect= −0.0381, p=0.542, N=419 across 35 lines). d, Spontaneous network activity was primarily driven by glutamatergic synaptic transmission as the application of glutamatergic inhibitors dl-AP5 (100µM) and DNQX (10µM) results in a 70.2% decrease in the percentage of active ROIs in both CON and SCZ lines (N=14 CON, 8 SCZ lines, p=2.15E-22). **e,** Representative traces of spontaneous excitatory postsynaptic currents. **f,** Frequency of sEPSCs are not significantly different between CON and SCZ (SCZ effect=-0.147, p=0.188, N=954 across 28 genomes). **g,** Amplitude of sEPSCs are not significantly different between CON and SCZ (SCZ effect=0.294, p=0.408, N=806 across 28 genomes).

**Extended Data Figure 7.**
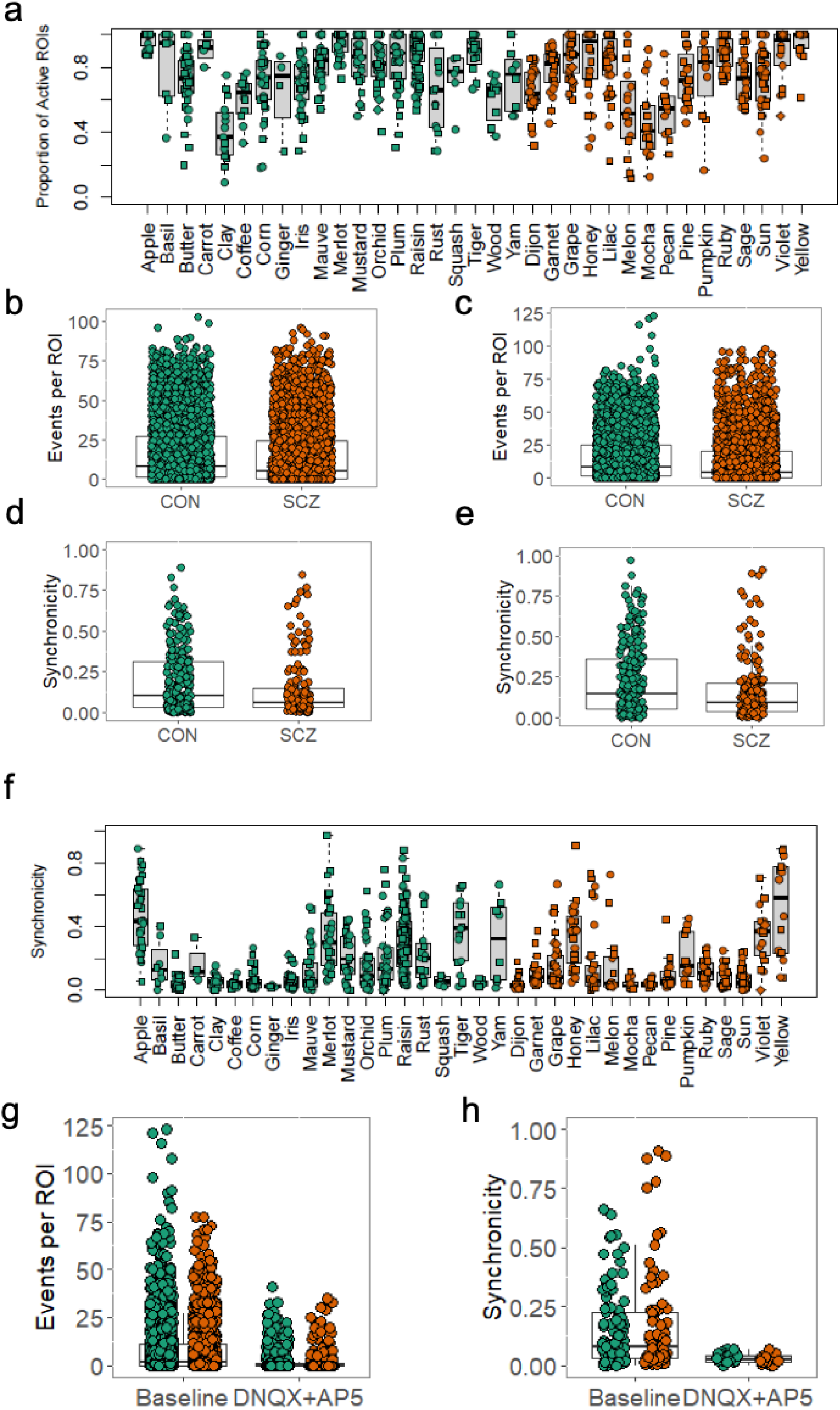
Summary data of calcium imaging by diagnosis and by line. **a,** Proportion of active ROIs by line (circle DIV42, square DIV63). **b,** Summary data of events per ROI by diagnosis at DIV42 (SCZ effect= −0.0136, p=0.9414, N_CON_=3902, N_SCZ_=2927). **c,** Summary data of events per ROI by diagnosis at DIV63 (SCZ effect=-0.040, p=0.828 N_CON_=3616, N_SCZ_=2866) **d,** Summary data of synchronicity of events by diagnosis at DIV42 (SCZ effect=-0.00383, p=0.9428, N_CON_=3902, N_SCZ_=2927). **e,** Summary data of synchronicity by diagnosis at DIV63 (SCZ effect=-0.0165, p=0.761, N_CON_=3616, N_SCZ_=2866). **f,** Synchronicity by line. **g,** Number of events per ROI is reduced following application of DNQX (10µM) and dl-AP5 (100µM), (baseline: N=2842, 8.37 ± 0.257, DNQX + AP5: N=739, 2.14 ± 0.204, p=1.34E-6) **h,** Synchronicity of events is decreased following application of DNQX (10µM) and dl-AP5 (100µM), (baseline: N=177, 0.160 ± 0.012, DNQX + AP5: N=38, 0.029 ± 0.00489, p=3.7E-5).

**Extended Data Figure 8.**
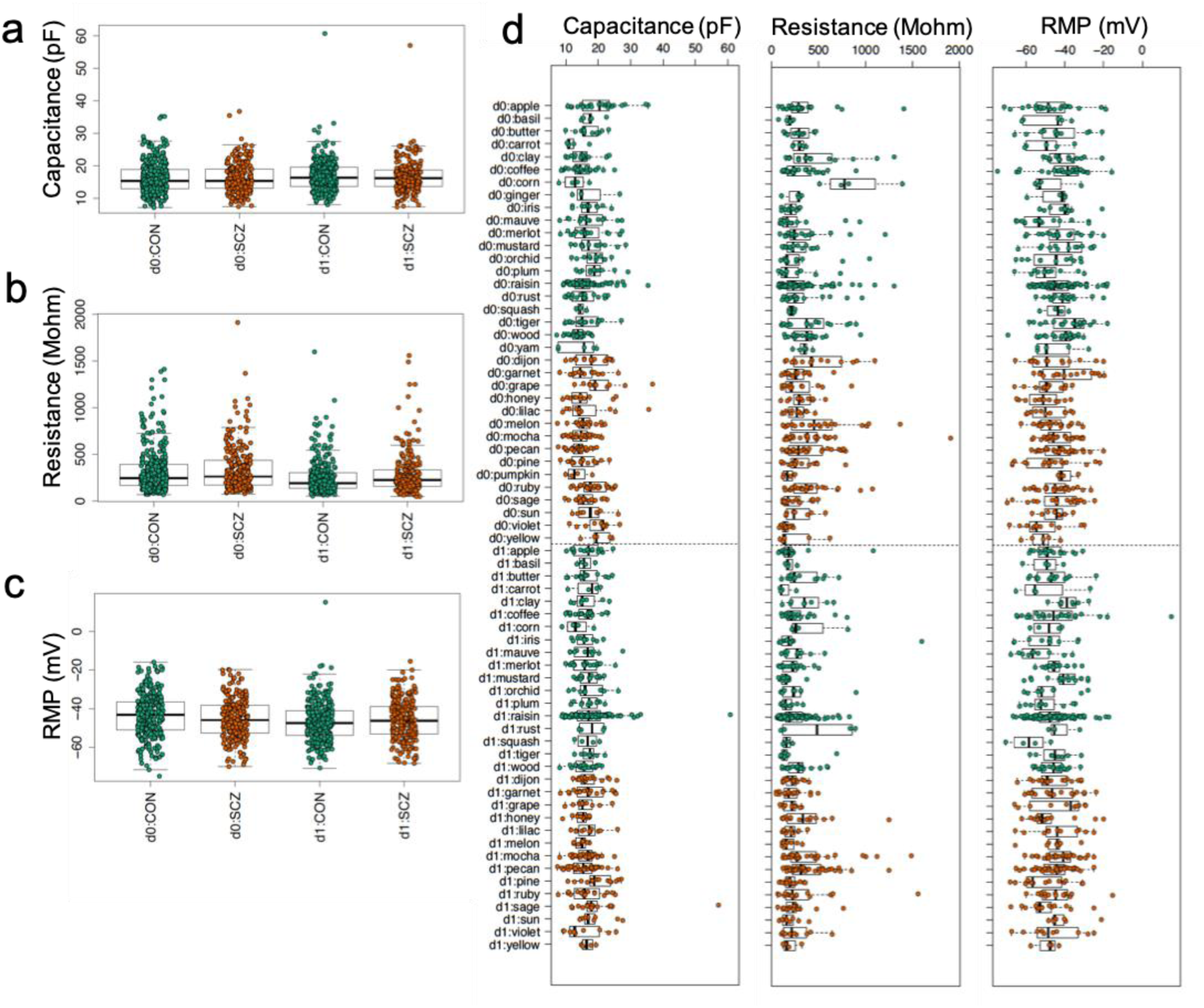
Neuronal maturation of membrane properties. **a,** Summary data by diagnosis showing membrane capacitance increases between DIV 56 (d0) and 70 (d1)(d1-d0 effect=0.799, p=0.003, N=1086), but is not different by diagnosis (Both timepoints SCZ effect=-0.043, p=0.878, N=1086; d0 SCZ effect=0.064, p=0.862, N=566 across 28 genomes; d1 SCZ effect=-0.182, p=0.666, N=520 across 28 genomes). **b,** Group data showing membrane resistance increases between DIV 57 (d0) and 70 (d1)(d1-d0 effect=-55.11, p=2.55E-05, N=1086), and is different by diagnosis (Both timepoints SCZ effect=29.41, p=0.026, N=1086; d0 SCZ effect=21.85, p=0.260, N=566 across 28 genomes; d1 SCZ effect=40.67, p=0.02, N=520 across 28 genomes). **c,** Summary data showing resting membrane potentials become more depolarized between DIV 57 (d0) and 70 (d1)(d1-d0 effect=-1.94, p=0.002, N=1086), but do not differ by diagnosis (Both timepoints SCZ effect=-0.665, p=0.295, N=1082; d0 SCZ effect=-2.022, p=0.023, N=565 across 28 genomes; d1 SCZ effect=0.890, p=0.331, N=517 across 28 genomes). **d,** Membrane property measures by line.

**Extended Data Figure 9.**
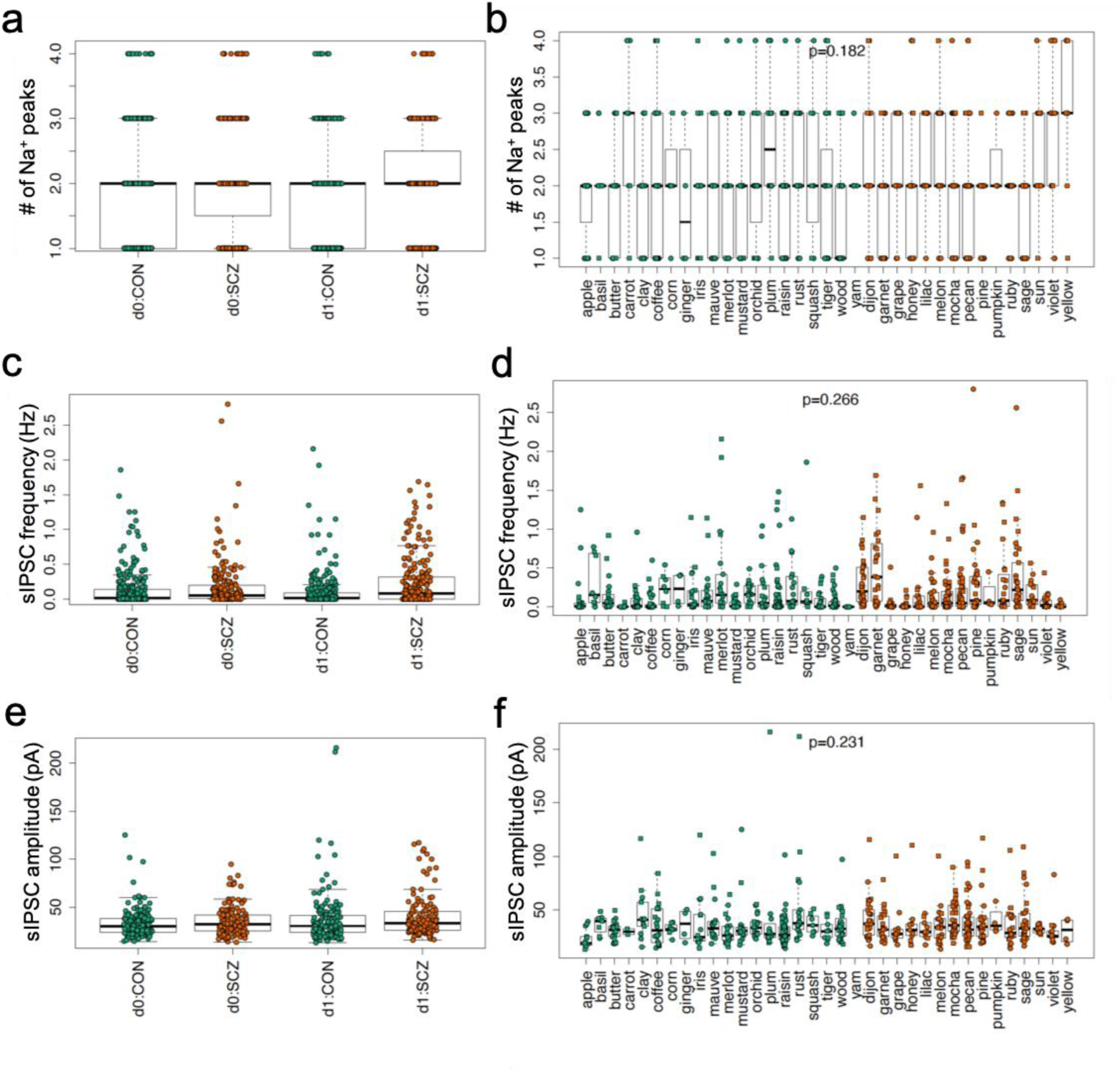
Summary data of number of Na^+^ peaks and sIPSC frequency and amplitude. **a,** Summary data by diagnosis showing number of Na^+^ peaks for DIV 56 (d0) and 70 (d1)(Both timepoints SCZ effect=0.093, p=0.038, N=1074 across 28 genomes; d0 SCZ effect=0.043, p=0.492, N=560 across 28 genomes; d1 SCZ effect=0.146, p=0.018, N=514 across 28 genomes). **b,** Summary data of Na^+^ peak counts for each line. **c,** Summary data by diagnosis showing sIPSC frequency for DIV 56 (d0) and 70 (d1)(Both timepoints SCZ effect=0.097, p=1.11E-06, N=903 across 28 genomes; d0 SCZ effect=0.049, p=0.082, N=449 across 28 genomes; d1 SCZ effect=0.143, p=2.10E-07, N=454 across 28 genomes).d, Summary data of sIPSC frequency for each line. **e,** Summary data by diagnosis showing sIPSC amplitude for DIV 56 (d0) and 70 (d1)(Both timepoints SCZ effect=2.131, p=10.184, N=591 across 28 genomes; d0 SCZ effect=1.903, p=0.250, N=304 across 28 genomes; d1 SCZ effect=2.421, p=0.350, N=287 across 28 genomes). **f,** Summary data of sIPSC amplitude for each line.

**Extended Data Figure 10.**
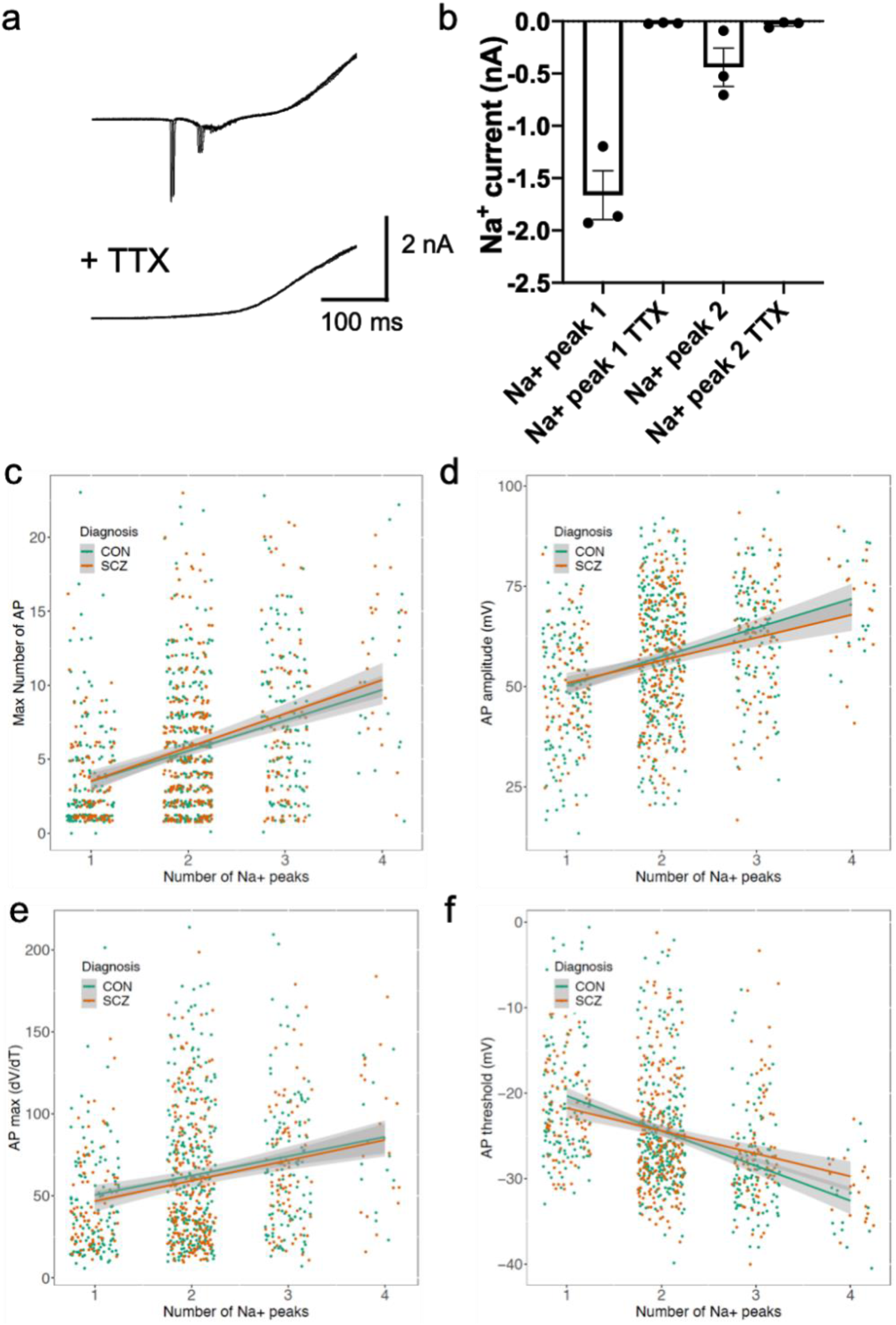
Physiological relevance of multiple Na^+^ peaks. **a,** Representative traces showing TTX blockade of Na^+^ currents in response to a voltage ramp. **b,** Group data showing TTX blockade of Na^+^ currents (peak 1 TTX effect=0.989, peak 2 TTX effect= 0.931, P_anova_=0.015, N=3). **c,** The maximum number of action potential is correlated with the number of Na^+^ peaks (CON N=625, R=0.347, p=1.87E-18; SCZ N=461, R=0.362, p=2.44E-15). **d,** The amplitude of action potentials are correlated with the number of Na^+^ current peaks (CON N=625, R=0.341, p=1.78E-15; SCZ N=461, R=0.279, p=1.65E-08). **e,** The acceleration of the action potential upslope (dV/dT) is correlated with the number of Na^+^ peaks (CON N=625, R=0.226, p=2.23E-07; SCZ N=461, R=0.245, p=7.89E-07). **f,** The action potential threshold is negatively correlated with the number of Na^+^ peaks (CON N=625, R=-0.430, p=1.67E-24; SCZ N=461, R=-0.296, p=2.00E-09).

**Extended Data Figure 11.**
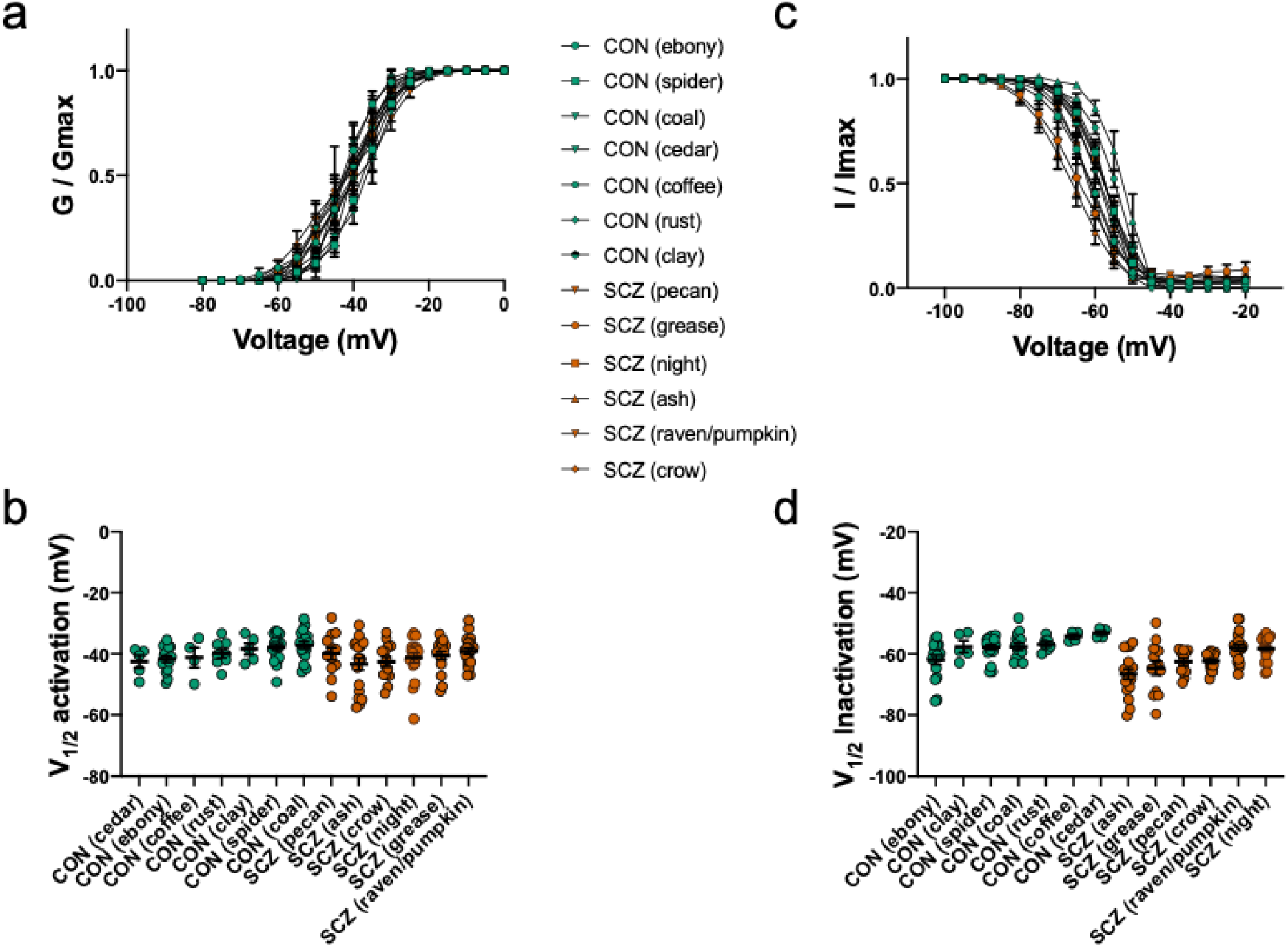
Summary data for Na^+^ channel activation and inactivation by genome. **a,** Normalized conductance versus voltage plots for 6 SCZ and 7 CON lines. **b,** Curves in (a) were fitted with a Boltzmann function and the V1/2 for each neuron is reported. **c,** Normalized inactivation versus voltage plots for 6 SCZ and 7 CON lines. **d,** Curves in (c) were fitted with a Boltzmann function and the V1/2 for inactivation for each neuron is reported.

## Acknowledgements

We are grateful for the vision and generosity of the Lieber and Maltz families, who made this work possible. This project was supported by the Lieber Institute for Brain Development. The collection of the clinical and cognitive data, and fibroblasts was supported by direct funding from the Intramural Research Program (IRP) of the NIMH to the Clinical Brain Disorders Branch (D.R.W., PI, protocol 95-M-0150, NCT00001486, annual report number ZIA MH002942-05) with supplemental support from the Clinical and Translational Neuroscience Branch (K.F.B., PI). We thank all of the participants in the IRP study and their families. We thank Flora Vaccarino for helpful advice and review of the manuscript, and Pat Levitt for his insights about the data.

## Author Contributions

D.R.W., B.J.M., and R.E.S. conceived and designed the study. S.C.P., S.R.S., Y.W., E.A.P., C.V.N., R.L.M., and O.R.S. collected RNA samples and performed RT-qPCR. S.R.S., Y.W., D.J.H., R.L.M. and O.R.S. performed stem cell culture and quality control experiments and neuronal differentiation. S.C.P., E.A.P., and C.V.N., performed immunocytochemistry and Ca2+ imaging experiments of neurons. M.N.T., N.J.E., J.M.S., and A.E.J. performed RNA-seq analysis. F.F., Z.Y. and H.Y.C. performed electrophysiology experiments. R.E.S. and A.E.J. carried out analyses of association between cellular measures and clinical/cognitive data. S.C.P., N.J.E, J.M.S., M.T., J.L.C and A.E.J. carried out data analyses. D.D., K.F.B., J.A.A., and D.R.W., provided clinical samples and clinical/cognitive data. S.C.P., S.R.S., K.M., A.E.J., R.E.S. and B.J.M wrote the manuscript. S.C.P., S.R.S., D.R.W., K.M., A.E.J., R.E.S., and B.J.M. edited the manuscript. D.R.W., K.M., A.E.J., R.E.S., and B.J.M supervised the study.

